# Cell generation dynamics underlying naïve T cell homeostasis in adult humans

**DOI:** 10.1101/635995

**Authors:** Jeff E. Mold, Pedro Réu, Axel Olin, Samuel Bernard, Jakob Michaëlsson, Sanket Rane, Andrew Yates, Azadeh Khosravi, Mehran Salehpour, Göran Possnert, Petter Brodin, Jonas Frisén

## Abstract

Thymic involution and proliferation of naive T cells both contribute to shaping the naive T cell repertoire as humans age, but a clear understanding of the roles of each throughout a human lifespan has been difficult to determine. By measuring nuclear bomb test-derived ^14^C in genomic DNA we determined the turnover rates of CD4^+^ and CD8^+^ naïve T cell populations and defined their dynamics in healthy individuals ranging from 20-65 years of age. We demonstrate that naïve T cell generation decreases with age, and that this could be explained by a combination of declining cell loss, peripheral division and thymic production during adulthood. We investigated putative mechanisms underlying age-related changes in homeostatic regulation of naïve T cell turnover using mass cytometry to profile candidate signaling pathways involved in T cell activation and proliferation in CD4^+^ naive T cells relative to CD31 expression, a marker of thymic proximity. We show that basal NF-κB phosphorylation inversely correlated with CD31 expression and thus is decreased in peripherally expanded naive T cell clones. Functionally we found that NF-κB signaling was essential for naive T cell proliferation to the homeostatic growth factor IL-7, and reduced NF-κB phosphorylation in CD4^+^CD31^−^ naive T cells is linked to reduced homeostatic proliferation potential. Our results reveal an age-related decline in naïve T cell turnover as a putative regulator of naïve T cell diversity and identify a molecular pathway that restricts proliferation of peripherally expanded naive T cell clones that accumulate with age.

## INTRODUCTION

Naïve T cell numbers and clonal diversity represent the adaptive immune system’s potential to sense and respond to foreign pathogens and mutant proteins expressed by malignant cells [1, 2]. In humans, the naïve T cell pool is established primarily in the first decade of life through the massive efflux of billions of newly produced naïve T cells from the thymus [3–5]. Each new naïve T cell is uniquely defined by a clonally heritable T cell receptor, which confers specificity for a restricted set of peptides for which a T cell is capable of sensing and responding against [6]. Thus, the number of unique foreign peptides that the T cell pool can sense is proportional to the number of unique clones present in the naïve T cell population at any given time.

It has long been recognized that the thymus undergoes a dramatic involution between birth and early adulthood in humans [4, 7]. Loss of thymic output in adults, shifts the responsibility for the maintenance of naïve T cell numbers to peripheral division of existing clones, rather than *de novo* production of new, unique clones [8]. In theory, this should lead to a gradual loss of naïve T cell diversity, as individual clones compete for space and limited homeostatic growth factors, such as IL-7, in the peripheral lymphoid tissues [9, 10]. Surprisingly, diversity and naïve T cell numbers are largely maintained until roughly 65 years of age in most humans, with only CD8^+^ naïve T cells showing an age-related decline in numbers during this period [10–13]. Notably, a dramatic loss of diversity has been observed in a fraction of elderly individuals, which has led to speculation that a sudden collapse of naïve T cell diversity may be causally linked to immune dysfunction associated with advanced age [10, 11].

Naïve T cells appear to divide rarely relative to other hematopoietic lineages, offering a putative mechanism for the maintenance of diversity over long periods of time [14]. Because naïve T cells divide infrequently, defining their actual turnover rates has been challenging, with estimates ranging from months to decades in different studies based on short-term pulse-chase labeling studies [15]. A major confounding feature of these studies is that only a very small fraction of the dividing naïve T cell population is labeled in the uptake period, and often no dilution of the label is observed in the short chase periods during which each human volunteer can be monitored [14, 16]. Additionally, recent evidence points towards greater heterogeneity within the classically defined ‘naïve’ T cell compartment, with some cell types present in this population exhibiting higher turnover than others, potentially leading to more rapidly dividing non-naïve cells being labeled and measured in these studies [1, 17]. Heterogeneity within truly naïve T cell populations is also known to exist. The markers CD31 and CD103 can subdivide CD4^+^ and CD8^+^ naïve T cells, respectively, into fractions with distinct replication histories defined by variable content of T cell receptor excision circles (TRECs) which are formed during naïve T cell development and diluted upon peripheral divisions [18, 19]. However, interpretation of these findings is complicated by the fact that differing TREC content may reflect thymic production of new T cells or selective outgrowth of certain clones in the periphery [20–22]. This also underscores the importance of combining TREC content with accurate cell age measurements when determining thymic activity in adult humans [23–25].

To better define the turnover rates of naïve T cells in healthy adult humans, we characterized the average cell age of millions of sort purified naïve T cells isolated from 59 healthy adults between 20 and 64 years of age by retrospective ^14^C dating of DNA [26]. Retrospective birth dating takes advantage of the large increase in atmospheric ^14^C due to nuclear bomb tests during the Cold War [27]. The ^14^C generated by the nuclear detonations reacted with oxygen to form ^14^CO_2_, which is taken up by plants through photosynthesis. The atmospheric ^14^C levels are mirrored in the human body at any given time, as we eat plants or animals that consume plants. Dividing cells incorporate ^14^C with a concentration corresponding to atmospheric ^14^C levels, creating a stable date stamp in the genomic DNA, which can be used to assess the age of cells and calculate turnover dynamics (Fig 1A). In this way, changes in ^14^C levels in the environment, and consequently the DNA of newly produced cells, function as a lifelong pulse chase experiment impacting all cells of an organism. This strategy has been successfully used to measure the rates of turnover of long-lived cells in humans including neurons, oligodendrocytes, adipocytes, cardiomyocytes and plasma cells [26, 28–31].

**Figure 1.**
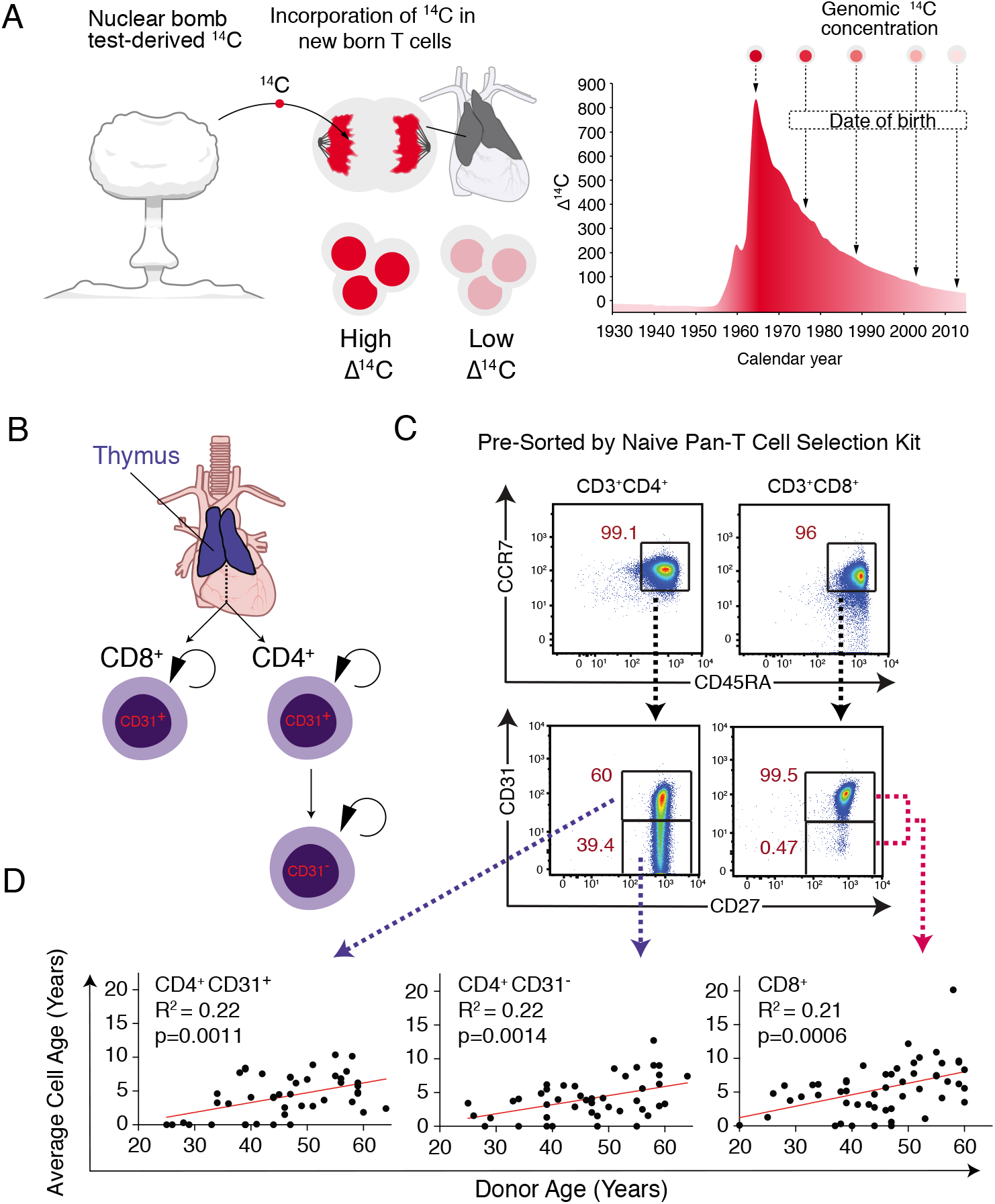
^14^C Measurements of Naïve T cell Populations Reveal Age-Dependent Decrease in Naïve T cell Turnover. (A) Schematic explaining how nuclear bomb test-derived ^14^C is incorporated into the DNA of newly produced naïve T cells (either in the thymus or through cell division in the periphery). Depending on what year a new naïve T cell is formed the ^14^C content will mirror atmospheric levels providing a cellular birthdate. (B) Populations of naïve T cells surveyed in this study include total CD8^+^ T cells and both CD4^+^CD31^+^ and CD31^−^ T cell populations. CD8^+^ and CD4^+^CD31^+^ naïve T cells are presumed to originate in the thymus and divide in the periphery with CD4^+^CD31^+^ cells yielding CD4^+^CD31^−^ cells in the peripheral tissues. (C) Flow cytometry plots depicting gating strategies for purifying each population of naïve T cells. Total CD3^+^ cells are shown for each gate which were obtained after purifying all naïve T cells from buffy coats subjected to red blood cell lysis. CD8^+^ naïve T cells are >98% CD31^+^ so no distinction was made for sorting this population. (D) Average cell age determined by ^14^C content analysis depicted for each population relative to donor age. Increased cell age is observed for all populations relative to donor age (student’s t-test).

Combining our cell age determinations with reported changes in peripheral naïve T cell number and TREC content, we provide a complete overview of how thymic activity, homeostatic division rates and naive T cell loss rates shape the peripheral naïve T cell pool throughout adulthood. We show for the first time, that the entire naïve T cell pool undergoes constant turnover in the periphery in healthy adult humans. We observed that turnover rates for both CD4^+^ and CD8^+^ naïve T cells decreased relative to donor age, consistent with recent predictions from mouse models [32]. Mathematical modeling shows that this decreased turnover was due to gradually reduced cell loss and peripheral division rates of naïve T cells with aging. For CD4^+^ naive T cells, we determined the average cell age for purified CD4^+^CD31^+^ recent thymic emigrants and CD4^+^CD31^−^ peripherally expanded naïve T cell populations. Modeling results indicated that CD31^−^ naive T cells may be limited in their ability to undergo additional homeostatic proliferation, while CD31^+^ naive T cells undergo peripheral homeostatic proliferation. This hypothesis was confirmed both in vivo and in vitro, and we provide a molecular mechanism based on differences in NF-κB phosphorylation relative to CD31 expression, that can account for this observation. Such a mechanism may in part explain how clonal diversity is preserved despite peripheral clonal competition for resources throughout decades of adult life. Our findings provide the first direct measurements of naive T cell lifespans throughout adulthood in humans and identify a potential mechanism that may in part explain the slow, regulated turnover of this population in the peripheral tissues.

## RESULTS

### Total CD4^+^ and CD8^+^ Naïve T Cell Turnover Rates Decrease Relative to Donor Age

In order to obtain accurate ^14^C measurements we required at least 3.5 million purified naïve T cells. CD4^+^ naïve T cells are typically more numerous in the blood and the fraction of CD31^+^ and CD31^−^ cells within this population typically afforded us the ability to obtain sufficient cell numbers for each. CD8^+^ naïve T cells were isolated as a single population due to cell number constraints (Fig 1B, C). A two-step purification strategy was developed to circumvent density centrifugation steps, required to remove granulocytes and red blood cells, which introduced contamination to downstream ^14^C measurements (data not shown). After red blood cell lysis, total white blood cell samples were subjected to a pan-naïve T cell isolation by magnetic beads (which also removes CD25^+^ naïve regulatory T cells) followed by FACS purification of CD45RA^+^CCR7^+^ CD4^+^CD31^+^, CD31^−^ or CD8^+^ T cells (Fig 1C). For CD8^+^ naïve T cells we included a subset of donors where we purified CD45RA^+^CD62L^+^CD3^+^CD8^+^ cells, the cell ages for these donors were not statistically different from those determined using our more stringent purification protocol based on CCR7 expression (S Fig 1).

We measured the ^14^C concentration in DNA isolated from purified naïve T cells from each population for all donors when possible, by accelerator mass spectrometry (Fig 1D, S Fig 1, S1 Table). The ^14^C content of a cell reflects both the ^14^C content of the 50% of its DNA that is inherited directly from its parent and the ^14^C content of newly synthesized DNA, which is assumed to reflect current atmospheric levels [26]. Despite the complexity of this process, it can be shown that the ^14^C content of a pool of cells at steady state, reflects the average time since a cell was produced, whether it be from the thymus or through homeostatic division, since the addition of a new cell is always associated with a complete new set of DNA (A detailed supplemental guide to all mathematical modeling is included entitled ‘Supplemental Mathematical Models (SMM))[26]. In simple terms, for a population at steady state, this average cell age is the inverse of the rate of turnover and is also the mean lifespan of all cells in the population.

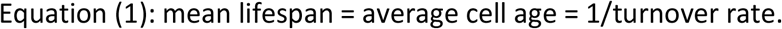

At steady state, the balance of loss and production of new cells from the thymus or by peripheral division, preserves the distribution of cell ages within the population. If the timescale over which a population undergoes turnover is short compared to the individual’s lifespan, we can use this cell age distribution together with the time course of recent levels of atmospheric carbon to estimate the mean of this distribution and hence the average lifespan of naïve T cells in an individual.

Stratifying these estimates by donor age, we observed that the mean age of CD4^+^CD31^+^, CD4^+^CD31^−^ and CD8^+^ naïve T cells all increased relative to donor age (Figure 1D, linear regression vs donor age: all p < 0.003). For all naïve T cells, the mean lifespans were of the order of 0-6 years (mean: 1.7yrs, sd: 2.1yrs) in 20-30 year olds, and 0-19 (mean: 6.6yrs, sd: 3.4yrs) in 50-65 year olds. This corresponds to an average annual turnover rate of 59% in 20-30 year olds and 15% in 50-65 year old donors (t-test, p<1e-6) (Fig 1D, S1 Table). We observed no statistically significant differences in naïve T cell age with respect to donor’s gender (data not shown).

### The Entire Naïve T Cell Pool Undergoes Constant Turnover Throughout Life

The heterogeneous nature of the naïve population [1] complicates the quantification of its homeostatic dynamics. In addition to phenotypically distinct cell types occupying the classical naïve T cell population, individual clones may exhibit differences in longevity or their propensity to respond to homeostatic growth signals [33, 34]. Dilution of TRECs across multiple decades of adult life could easily result from uneven clonal expansions, where only a fraction of the naïve T cell pool is selectively expanding in the periphery [20]. Likewise, short-term labeling studies only label a small percent of the total naïve T cell population (0-5%), meaning that no inference can be made concerning the potential of the remaining cells to undergo division [14, 16].

By measuring the average age of millions of naïve T cells from donors born at various times before, during and after the atmospheric ^14^C bomb-spike, we can address whether the naïve T cell compartment contains subpopulations of long-lived non-dividing cells. If a large fraction of naïve T cells were to be produced during the first decades of life and would recirculate without undergoing subsequent homeostatic division, these cells would contain ^14^C levels that match the environment from an individual’s childhood. This would result in donors born during the peak of the bomb-spike to have elevated ^14^C levels as a result of accumulated non-dividing naïve T cells generated during this timeframe.

We compared one model in which all cells are equally likely to divide (A) with a second model (2POPA) where populations of different sizes accumulate in the periphery without undergoing additional rounds of division (Fig 2, SMM p.12 and S2 Table). Although our samples show increasing average cell age relative to donor age (Fig 1D), the best fit for our data indicates a linear increase in cell age, whereas an exponential increase would indicate the presence of non-dividing cells generated early in life (Fig 2). Our data therefore suggest that the majority (99%) of naïve T cells are maintained dynamically throughout life through the combination of the production of new cells by the thymus, proliferative renewal of existing cells, and loss through death or differentiation.

**Figure 2.**
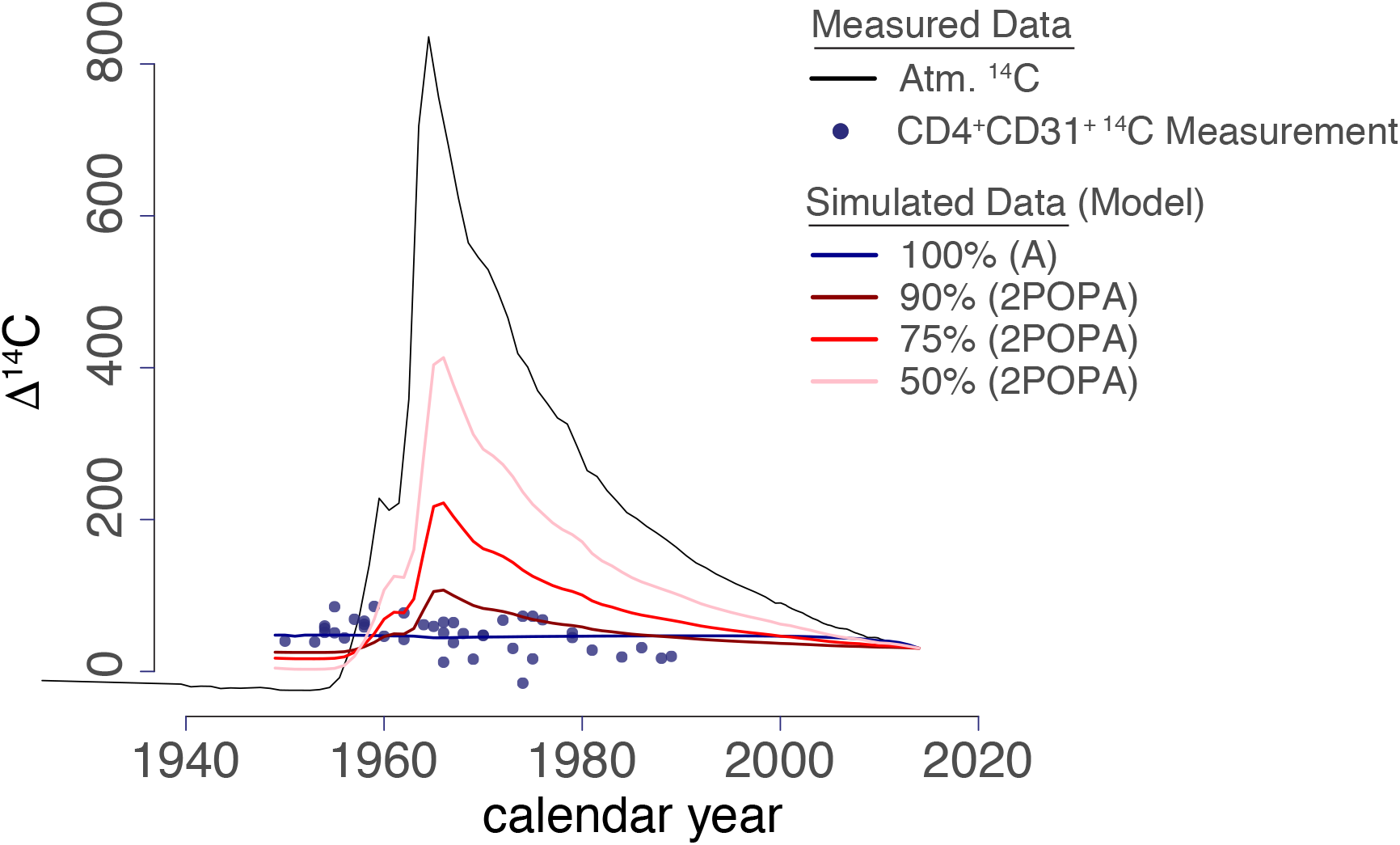
All Naïve T Cells Undergo Turnover Throughout Adult Life. Individual data points for ^14^C measurements of DNA isolated from CD4^+^CD31^+^ naïve T cells plotted according to donor birthyear (x-axis). The atmospheric ^14^C levels are plotted (black line) and simulated data are shown to represent the average distributions of datapoints if 50%, 75%, 90% or 100% of the naïve T cells undergo turnover after production. If a fraction of the naïve T cell population persisted as non-dividing cells in the periphery after thymic egress, donors born between 1960-1980 would have elevated ^14^C content measured in the DNA of the naïve T cell population. Model testing was performed for CD4^+^CD31^+^, CD4^+^CD31^−^, total CD4^+^ naïve, and total CD8^+^ naïve T cell populations and all results were consistent with 100% turnover (Table S2).

### Modeling Thymic Output, Peripheral Division, and Cell Loss Rates Based on Cell Age Estimates

Our estimates of cell age allow us the opportunity to define the dynamic turnover rates for naïve T cells throughout an adult human lifetime. To do this, we incorporated regressed data taken from publications for both CD4^+^ and CD8^+^ naïve T cells relating to cell numbers [13], T cell receptor excision circles (TREC) [8, 23], and cell age estimates defined by ^14^C measurements (S Fig 2). Individually, each of these measurements gives insight into naïve T cell turnover throughout life. Healthy humans ranging from 20-65 years of age show minimal decline in CD4^+^ naïve T cell numbers while CD8^+^ naïve T cell numbers progressively decline in this timeframe [13]. TREC content measurements in sorted naïve T cells allow insights into cell loss and proliferation [24]. Finally, total cell age measurements allow us to demonstrate that the entire naïve T cell pool is subject to division and define the average rate of division for any naïve T cell. Using a series of linear equations, similar to those previously used to define T cell dynamics during the first decades of human life [5, 35], we set out to quantify the processes that impact the maintenance of naïve T cell numbers and diversity throughout adult life.

For naïve CD8^+^ T cells there were four observables for each donor: (1) cell number, (2) TREC content, (3) cell age and (4) CD31 expression, which does not undergo age-related downregulation and thus was set to 98% which was the average frequency in our dataset. We therefore had four kinetic parameters to solve for: thymic export rate (F), peripheral proliferation rate *(rho),* loss rate *(gamma),* and a small CD31 marker loss rate (described in SMM). It was assumed that the CD31^+^ and CD31^−^ subsets had the same age, and that the CD31^−^ T cells did not proliferate significantly (Fig 3A). Under these conditions, a unique kinetic parameter set could be found for each donor (Fig 3B).

**Figure 3.**
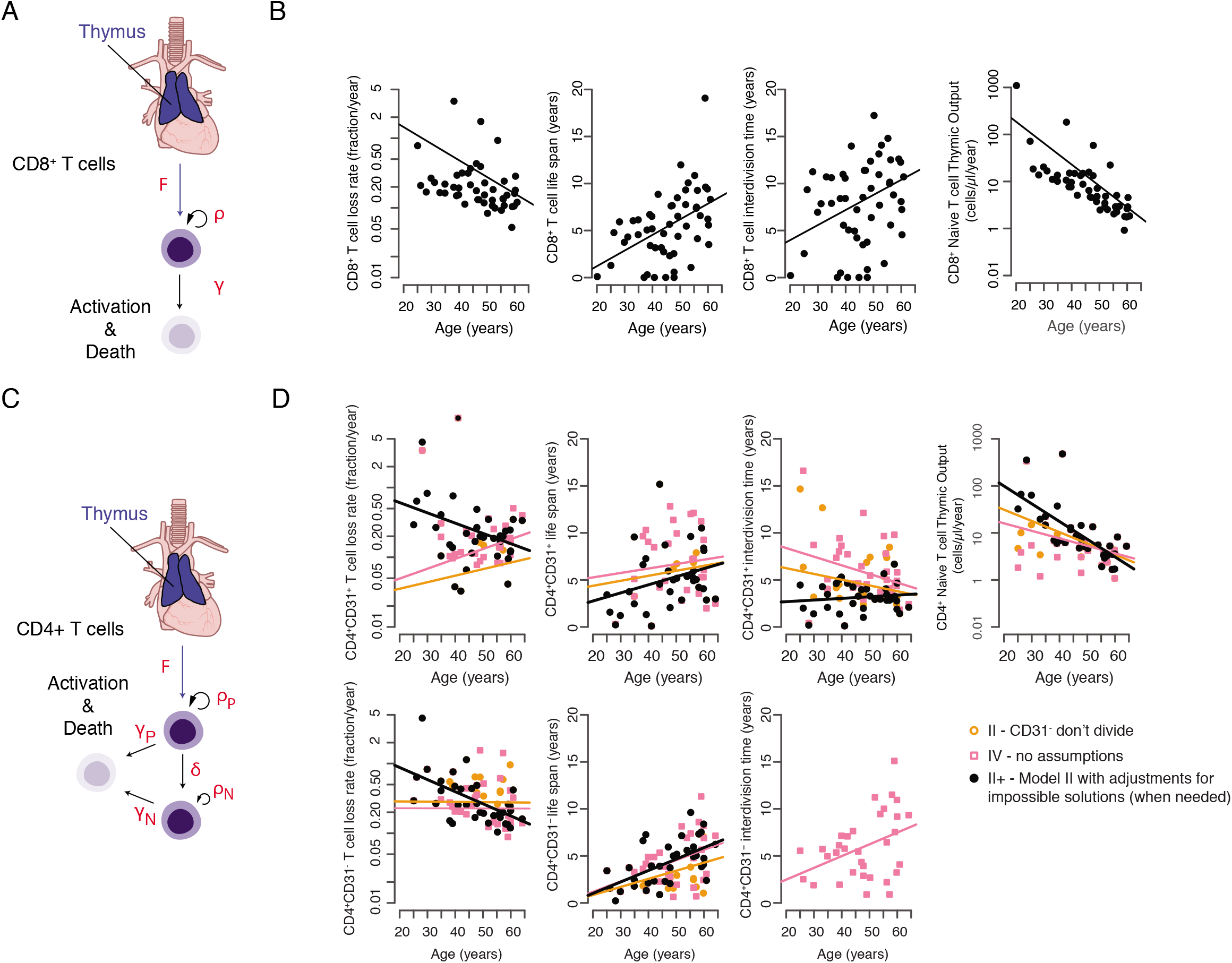
Dynamics of Naïve T cell Homeostasis Determined by Linear Models. (A) Individual variables measured with linear equations depicted for CD4^+^ naïve T cell populations. (B) Solutions for linear models are shown for selected scenarios tested (from Table S3). Scenario IV (pink squares) shows solutions when no predictions are made and each dynamic variable is solved with no additional assumptions. Scenarios II and II^+^ show solutions when CD31-naïve T cells are assumed to be a non-dividing population. Scenario II^+^ is modified to adjust for individuals where an exact solution could not be made (n=7/41 donors, SMM). (C) Individual variables measured with linear equations depicted for CD8^+^ naïve T cell populations. (D) Solutions for each dynamic variable with no additional assumptions made are depicted for CD8^+^ naïve T cells.

CD4+ naïve T cells had six observables and six dynamic variables to determine. Here we again used information regarding cell numbers, TREC content, and cell age, but defined individual-specific values for both CD4^+^CD31^+^ and CD4^+^CD31^−^ fractions as well as an additional dynamic variable to account for the differentiation of CD4^+^CD31^+^ cells to CD4^+^CD31^−^ (Fig 3C). Importantly, the processes governing loss of CD31 expression by naïve CD4^+^ T cells in humans remains unknown, although it is generally assumed that the CD31^−^ population accumulates due to expansion of CD31^+^ cells coupled with downregulation of CD31 expression with age [36]. We estimated the following variables for CD4^+^ naive T cells; The rate of export of CD4^+^CD31^+^ naïve T cells from the thymus, F; and the division rates of CD4^+^CD31^+^ and CD4^+^CD31^−^ T cells, *rho_P* and *rho_N,* respectively. Upon division, TREC content is split with equal probability between daughter cells. There is also a probability *delta* for CD31^+^ T cells to lose the CD31 marker, and to produce two CD31^−^ T cells. The two populations are lost though death or differentiation at rates *gamma_P* and *gamma_N.* The resulting model is expressed as an algebraic linear system of six equations and six kinetic parameters to solve for. For each individual, conditions for existence and uniqueness of positive parameters were defined, and an optimal set of kinetic parameters computed (Fig 3C and D, S Fig 4).

### Dynamics of Naïve T cell Turnover According to Modeling

Cell number can be maintained by thymic output, by peripheral proliferation, and/or lower cell loss. Our calculations suggest that total thymic output declines exponentially with age. Loss rates decline for both CD4^+^ and CD8^+^ naïve T cell populations relative to age, but did not differ significantly between cell types (Fig. 4A, B, Wilcoxon rank sum test, p>0.05). Proliferative activity was significantly higher in the CD4^+^ population (Wilcoxon rank sum test, p<1e-6). Although the net proliferative output in the CD4^+^ and CD8^+^ naïve T cell populations declines with donor age, the contribution to cell number maintenance increases, from around 60% at age 20 to more than 95% at age 65, owing to a substantial decline in thymic output over this timeframe. Total thymic output in the CD4^+^ naïve T cell population is twice that of the CD8^+^ naïve T cell population (Fig 4A, B, Wilcoxon rank sum test, p=0.002). However the relative contribution of thymic output to the peripheral CD8+ naïve T cell population is slightly higher, due to the numerical differences in peripheral proliferative output (Fig. 4A-C, Wilcoxon rank sum test, p=0.04).

**Figure 4.**
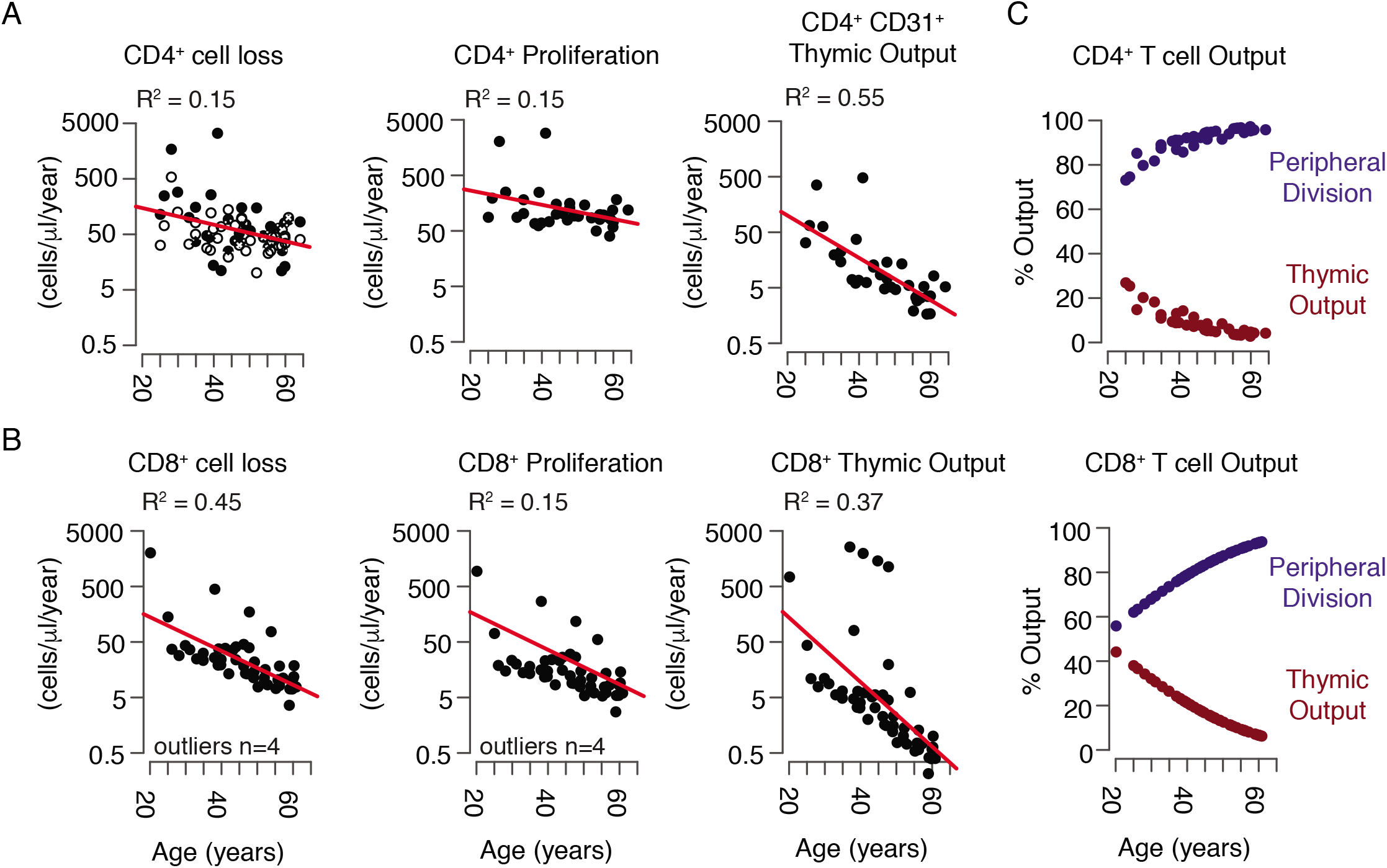
The CD4^+^ and CD8^+^ Naïve T cell Populations Exhibit Reduced Cell Loss, Peripheral Division, and Thymic Production Throughout Adulthood. (A) Dynamic rates of cell loss, homeostatic turnover and thymic activity for CD4^+^ naïve T cells calculated as a total population (combining information for CD4^+^CD31^+^ and CD31-cells for loss and proliferation rates). (B) Dynamics of CD8^+^ naïve T cell loss, proliferation and thymic activity relative to donor age. (C) Relative contribution of thymic activity and peripheral turnover for CD4^+^ (top) and CD8^+^ (bottom) naïve T cell populations expressed as %Output or fraction of new cells contributed by each mechanism.

A common explanation for how total CD4^+^ naïve T cell numbers persist throughout adulthood relative to CD8^+^ naïve T cells, rests primarily on their ability to expand as they differentiate into CD4^+^CD31^−^ naïve T cells [12, 36]. Our data on cell age provide the first estimate of the rate at which this transition occurs, as we see that CD4^+^CD31^+^ cells exhibit increased average cell ages in older donors (Fig 1D, 3D), indicating that CD31 may be better treated as a marker of limited CD4^+^ naïve T cell peripheral expansion rather than recent thymic egress.

### Modeling the Relationships between CD4^+^CD31^+^ and CD4^^+^^CD31^−^ Naïve T cells Suggests CD31^−^ T cells are Terminally Expanded

Little is known about the mechanisms leading to loss of CD31 expression, or the consequences of CD31 downregulation in the human CD4^+^ naïve T cell subset [37, 38]. It is clear that a similar process does not occur in CD8^+^ naïve T cells, or in mouse models. CD4^+^CD31^−^ naïve T cells accumulate throughout adult life, and exhibit decreased TREC content, potentially indicating that peripheral expansion of CD4^+^ naïve T cells coupled with loss of CD31 expression maintains a steady state in CD4^+^ naïve T cell numbers throughout adulthood [18, 38, 39]. As a population, CD8^+^ naïve T cells contain comparable TREC content to CD4^+^CD31^+^ naïve T cells, and the number of CD8^+^ naïve T cells declines with age [8, 13, 23].

Because CD4^+^CD31^−^ naïve T cells have decreased TREC content and accumulate over time, we anticipated that this population would have a higher rate of turnover and hence a shorter lifespan than CD4^+^CD31^+^ cells. Our results, however, indicate that the two populations age comparably. Moreover, the age-related increase in CD4^+^CD31^+^ naïve T cell lifespan suggests that CD31 expression is not indicative of recent thymic production, but rather reflects peripheral division history, as CD31^+^ naïve T cells in 50-64 year old individuals had average cell ages of 4-11 years. We therefore decided to test several hypotheses concerning the dynamic behavior of each population of CD4^+^ naïve T cells using our linear models (S Fig 4). In particular, we wanted to assess whether our data were consistent with the possibility that CD31^−^ naïve T cells represent a terminally expanded population of naïve CD4^+^ T cells which is constantly fed by homeostatically dividing CD4^+^CD31^+^ naïve T cells throughout life.

We addressed this hypothesis by setting the peripheral division rates of CD4^+^CD31^−^ naïve T cells to 0 in our linear models (H2). Additional hypotheses were that death rates of CD31^+^ and CD31^−^ cells were equal (H1) and CD31^+^ naïve T cells do not undergo homeostatic division (H3), in each case resulting in removal of a single dynamic variable (S Fig 4). We determined goodness of fit by measuring the sum of squared errors (SSE) and the differences in Akaike Information Criterion values (ΔAICc) as a measure of model accuracy [40]. Only the scenario in which CD31^−^ naïve T cells were assumed to be a non-dividing population (Scenario II) provided a better potential fit based on the ΔAICc than our model with no assumptions (Scenario IV), though each had roughly equivalent SSE (S Fig 4). Thus, our data are consistent with the hypothesis that CD4^+^CD31^−^ naïve T cells do not undergo peripheral homeostatic division. Notably, our initial model (scenario IV) indicated an age-related decline in CD31^−^ naïve T cell proliferation, but not CD31^+^ naïve T cell proliferation, consistent with the idea that CD31^+^ naïve T cells have heightened homeostatic proliferative potential.

### CD31 Expression on CD4^+^ Naïve T cells is Linked to Homoeostatic Proliferative Potential *In Vivo*

Two previous studies have reported that CD4^+^CD31^−^ naïve T cells may have limited proliferative potential to IL-7 *in vitro* [41, 42]. It remains unclear, however, whether differences exist for each population *in vivo,* and no attempts have been made, to our knowledge, to measure the proliferation of each population using markers that exclude contaminating memory populations in the naïve T cell pool. We therefore profiled 11 healthy adult donors (average age 48±16 years, range 25-69) using a comprehensive panel of markers specific to naïve T cells (CD4^+^CD45RA^+^CCR7^+^CD27^+^CD127^+^CD57^−^PD1^−^CD95^−^) and the proliferation marker Ki67, which labels cells that underwent mitosis within the preceding 72 hours [43], to define the frequency of dividing naïve T cells in relation to CD31 expression in vivo (Fig 5A-C). Using these stringent criteria to focus on truly naïve T cells, we observed there were very few naïve T cells exhibiting evidence of recent proliferation (0.003-0.031%) as expected (Fig 5B, C, E, and F). By contrast T stem cell memory (T_scm_) cells (CD4^+^CD45RA^+^CCR7^+^CD27^+^CD127^+^CD57^−^PD1^−^CD95^+^), which have a similar phenotype as CD4^+^ naïve T cells apart from expression of CD95, exhibited much higher frequencies of dividing cells, again indicating that this population may impact turnover estimates in prior studies (Fig 5D). We noted that CD31 expression within the CD4^+^ naïve T cell population is better viewed as a continuum rather than discreet CD31^+^ and CD31^−^ populations (Fig 5B). Therefore, we gated on CD31^bright^, CD31^dim^ and CD31^−^ populations and quantified Ki67 frequencies in each (Fig. 5B, E). We noted that the CD31^−^ population showed significantly less proliferation across all donors (0.00204%-0.023%, mean: 0.01%, s.d.: 0.006%) providing additional support for our hypothesis that these cells exhibit limited peripheral turnover potential (Fig 5E). CD8^+^ naïve T cells had high CD31 expression and comparable frequencies of dividing cells to CD4^+^CD31^+^ naïve T cells in all donors (range 0.0099%-0.024%, mean: 0. 017%, s.d.: 0.005%) (Fig 5C, F).

**Figure 5.**
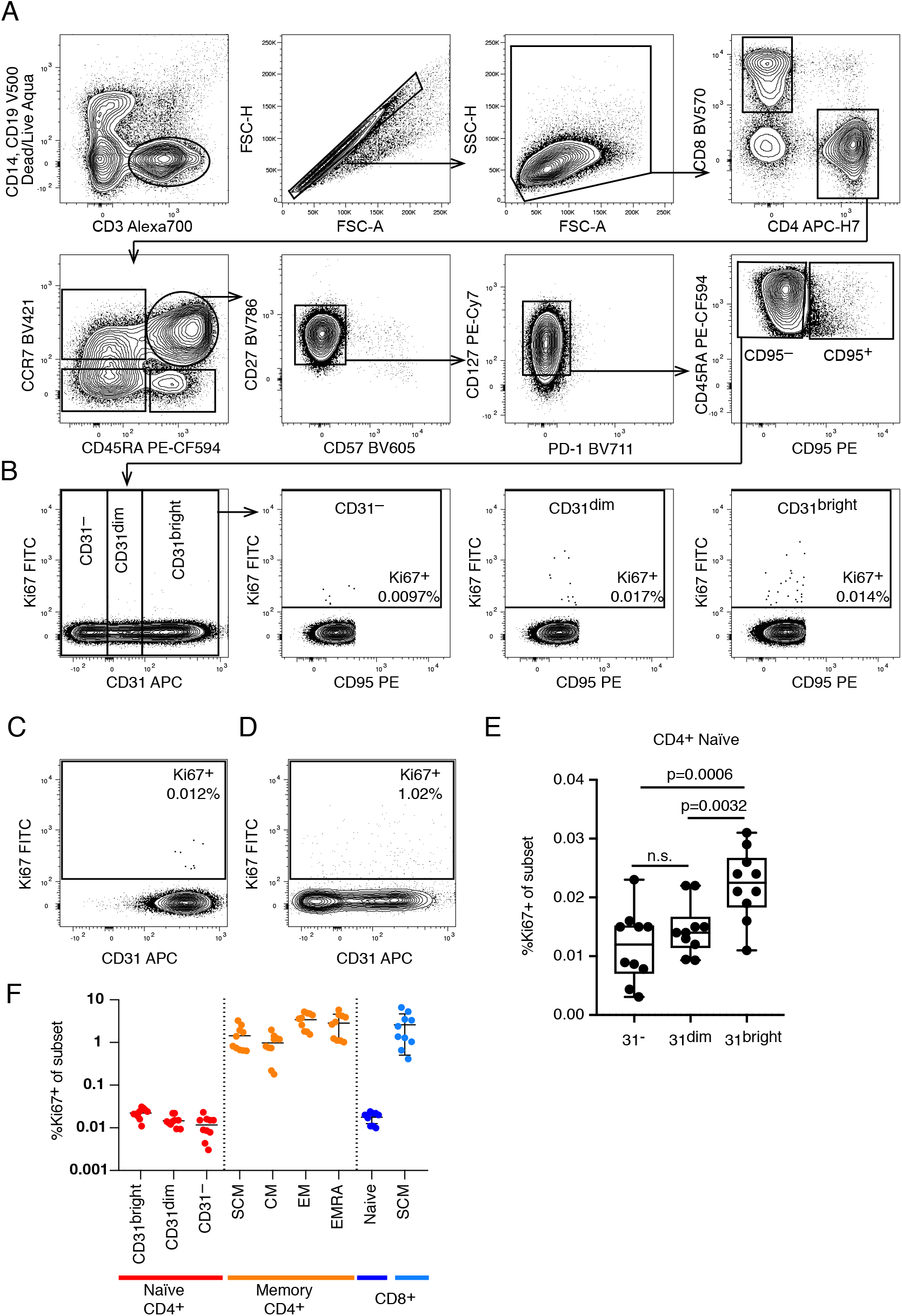
CD4^+^CD31^+^ Naïve T cells Proliferate more than CD4^+^CD31^−^ Naïve T cells In Vivo. (A) Gating strategy to identify naïve T cells. Naïve CD4^+^ T cells were identified as live, CD14^−^ CD19^−^CD3^+^CD4^+^CD8^−^CD45RA^+^CCR7^+^CD57^−^CD27^+^PD1^−^CD127^+^CD95^−^ cells to exclude potential contaminating populations of activated cells including stem cell memory cells. The same strategy was used to identify CD8^+^ naïve T cells by switching to the CD4^−^CD8^+^ fraction. (B) Representative example of CD4^+^ naïve T cell distribution for CD31 expression and gating strategy to define CD4^+^CD31^bright^, CD4^+^CD31^dim^, and CD4^+^CD31^−^ cells. (C) Frequency of Ki67^+^ dividing cells in the CD8^+^ naïve T cell gate. (D) Frequency of Ki67^+^ dividing cells within the stem cell memory population present in the conventional CD4^+^CD45RA^+^CCR7^+^ gate. (E) Summary of all donors showing frequency of Ki67^+^ dividing cells for each population identified with our gating strategy. Significantly less Ki67^+^ cells were observed in the CD4^+^CD31^−^ Naïve T cell fraction as compared to the CD31^bright^ population (p=0.0006, paired t test) and the CD31^dim^ vs CD31^bright^ population (p=0.0032, paired t test). (F) Frequency of Ki67^+^ cells in CD4^+^ Naïve T cells, CD4^+^ Memory T cell subsets, and in CD8^+^ Naïve and CD8^+^ stem cell memory subsets for all donors.

### Single Cell Proteomic Analysis of CD4^+^ Naïve T cells Reveals a Link Between CD31 Expression and Basal NF-κB Phosphorylation

We reasoned that we could potentially examine molecular pathways regulating homeostatic proliferation by examining CD4^+^ naïve T cells relative to CD31 expression. To this end, we utilized mass cytometry to interrogate differences in both phenotypic and functional properties of single naïve T cells that correlated with differences in CD31 expression level. This allowed us to more accurately address differences according to the range of CD31 expression detected within the total CD4^+^ naïve T cell pool (Fig 5B).

We first confirmed that CD45RA^+^CCR7^+^CD4^+^CD31^+^ and CD45RA^+^CCR7^+^CD4^+^CD31^−^ naïve T cells were phenotypically indistinguishable with respect to 14 additional T cell activation and differentiation markers including the homeostatic cytokine receptor IL-7R (S4 Table) visualized using t-stochastic neighborhood embedding (t-SNE) and ACCENSE (Figure 6A) [44, 45]. Next, we performed *in vitro* stimulation assays on total peripheral blood mononuclear cells and monitored signaling states of individual cells by mass cytometry with antibodies targeting phospho-proteins involved in T cell activation pathways (ERK1/2 and SLP-76) [46]. Temporal changes in the activation of different signaling pathways were directly related to differences in CD31 expression using standard Pearson’s correlations and a recent conditional-density estimator called DREVI (conditional-density rescaled visualization) [46]. With DREVI the density estimate of the response variable is renormalized to obtain a conditional-density estimate of this variable given the abundance of another variable, CD31 cell surface expression in this case. We observed the expected kinetic changes in phosphorylation of canonical signaling molecules SLP-76 and ERK upon T cell receptor crosslinking, but these patterns showed no relationship with CD31 expression (Figure 6B).

**Figure 6.**
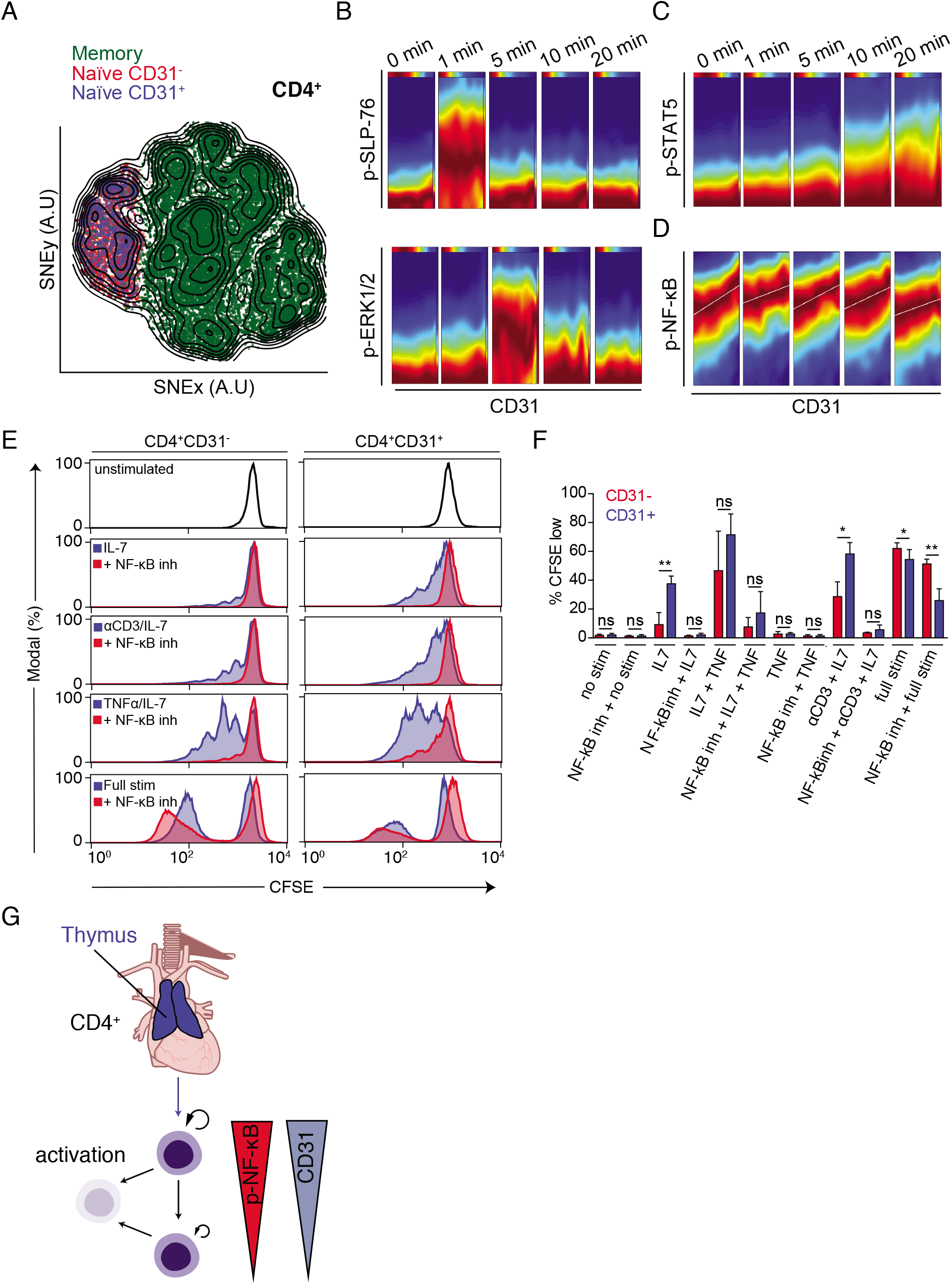
Evidence for a Link Between Replicative History and Homeostatic Proliferation Potential Controlled by NF-κB. (A) Memory/effector (green) and naïve CD4^+^ T cells divided into CD31^−^ (red) and CD31^+^ (blue) cells, were merged and their high-dimensional single-cell phenotypes analyzed by mass cytometry visualized using a tSNE dimensionality reduction. (B) DREVI plots depicting correlation between CD31 and the phosphorylation of SLP-76, ERK1/2, (C) STAT5 and (D) NF-κB throughout a 20-minute homeostatic stimulation with anti-CD3 and IL-7. Data are representative of 4 separate experiments on 8 donors. (E) Proliferation of sort purified CD4^+^CD31^−^ (left) and CD4^+^CD31^+^ (right) naïve T cells in response to different stimulation conditions (blue histograms) as well as in the presence of the the NF-κB inhibitor 6-amino-4–4-phenoxyphenylethylamino-quinazoline (red histograms). Data represent 4 experiments on 8 donors. (F) Summary data from four independent donors showing percent CFSE^low^ (proliferated) cells within the CD31^+^ and CD31^−^ fraction of naïve CD4^+^ T cells relative to different stimulatory conditions. Cells were treated with vehicle (DMSO) or NF-κB inhibitor (6-amino-4-4-phenoxyphenylethylamino-quinazoline) for each stimulation condition. Full Stim: αCD3/αCD28/IL-2. (G) Schematic summary of the dynamics of CD4^+^ naïve T cell homeostasis in adult humans.

We addressed whether differences in CD31 expression levels correlated with differential responses to the naïve T cell growth factor IL-7. Because TCR engagement has been implicated as an important component of homeostatic signaling of naive T cells in mice, we also included TCR crosslinking with antibodies against CD3 in our assays [47]. Notably, we observed that inclusion of CD3 crosslinking resulted in only a modest increase in naive T cell proliferation and evidence of a small degree of T cell activation as indicated by loss of CD45RA expression (S Fig 5A). The proximal signaling event downstream of IL-7R stimulation is the phosphorylation of STAT5. We observed that phosphorylation of STAT5 in response to IL-7 occurred within 10 minutes after stimulation and increased throughout the course of stimulation. STAT5 phosphorylation was found to be equivalent across the CD4^+^ naïve T cell population regardless of CD31 expression (Fig 6C). This observation was consistent with the fact that IL-7R levels were equivalent for naïve CD4^+^CD31^+^ and CD31^−^ cells (Fig 6A).

A comprehensive examination of all changes in measured phospho-protein expression relative to CD31 expression in naive CD4^+^ T cells revealed a consistent positive relationship between CD31 expression and phosphorylation of the NF-κB subunit p65 (RelA) (Fig 6D and S Fig 6, S4 Table). This relationship was observed in the absence of stimulation as well as in the presence of IL-7 and CD3 crosslinking. Stimulation with tumor necrosis factor (TNF), which is known to act through NF-κB, resulted in a slight increase in NF-κB phosphorylation across the naive CD4^+^ T cell population regardless of CD31 expression (S Fig 5B). Interestingly, basal NF-κB phosphorylation has been shown to play an important role in determining naïve T cell survival in rodents [48, 49], prompting us to investigate this pathway further.

### Homeostatic Proliferation in Response to IL-7 is Dependent Upon NF-κB Signaling

We next determined whether CD4^+^CD31^+^ and CD4^+^CD31^−^ Naive T cells showed differences in proliferative potential in vitro in response to IL-7 stimulation using CFSE dilution assays. Consistent with prior reports, we noted a significant reduction in proliferation in response to IL-7 stimulation by CD4^+^CD31^−^ naive T cells relative to CD4^+^CD31^+^ naive T cells (Fig 6E). Both populations proliferated and underwent differentiation in response to T cell activation stimuli, with CD4^+^CD31^−^ naive T cells exhibiting a small but statistically significant increase in proliferation. Interestingly, CD8^+^ naive T cells were highly responsive to IL-7 stimulation *in vitro* despite exhibiting generally lower turnover rates *in vivo* as measured by ^14^C levels.

To address whether differences in basal NF-κB phosphorylation levels account for differential proliferative potential of CD4^+^CD31^+^ and CD4^+^CD31^−^ naïve T cells, we measured homeostatic proliferation of each population in the presence of an NF-κB inhibitor (6-amino-4–4-phenoxyphenylethylamino-quinazoline). Inhibition of NF-κB completely abolished homeostatic proliferation, but did not completely inhibit proliferation in response to activation signals (Fig 6E, F) [41]. Additionally, we found that we could rescue homeostatic proliferation of the non-responsive CD4^+^CD31^−^ naïve T cells by promoting NF-κB phosphorylation through the addition of TNF (Fig 6E, F). Notably, proliferation in the presence of TNF was still dependent upon IL-7 and did not result in activation of naïve T cells as determined by CD45RA downregulation (Fig 6F, S Fig 5C). We observed a similar dependence on NF-κB activity for homeostatic proliferation of CD8^+^ naïve T cells, which in general showed heightened responses to IL-7 stimulation (S Fig 5B, C). Thus, our data support a model where gradual loss of CD31 expression by naïve CD4^+^ T cells is associated with decreased NF-κB phosphorylation leading to reduced peripheral homeostatic proliferative potential, but comparable activation potential (Fig 6G).

## DISCUSSION

Here we provide a comprehensive dataset defining the global turnover rates of the naïve T cell pool in adult human beings until 65 years of age. Our findings extend nicely upon estimates made concerning the dynamic regulation of the generation of the naïve T cell pool during the first 20 years of life [5]. Our conclusions were not wholly consistent, however, with a recent study tracking naïve T cell turnover in donors between 65-75 years of age, where no age-related changes in turnover were documented relative to younger donors [16]. This discrepancy may reflect the fact that individuals over 65 years of age contain elevated frequencies of memory cells and activated T cells within the population traditionally identified as naïve T cells, as well as changes that are reported to occur in advanced age that lead to collapse of naïve T cell diversity [11, 50, 51]. Additional efforts to define the cell age of healthy humans over 65 years of age will be needed to better address these differences.

It is now recognized that the human naïve T cell pool fundamentally differs from laboratory mice, with respect to factors responsible for maintaining cell numbers and diversity throughout life [8]. Mice maintained in pathogen free environments typically live for two to three years, throughout which time the thymus accounts for nearly all new production of naïve T cells [52]. In humans, the thymus is capable of generating hundreds of millions of new naïve T cells every year until roughly 30 years of age, as predicted by our models as well as previously published estimates [8]. Histological data support this conclusion, as medullary and cortical regions in which T cell development occurs, are visible until roughly 40 years of age [4, 7]. The loss of thymic function after 40 years has prompted speculation that sudden increases in human lifespans have outpaced evolutionary processes needed to maintain an adaptive immune system in older age [53]. Nonetheless we maintain a diverse naïve repertoire for several decades beyond the time that the thymus ceases to function, due primarily to peripheral homeostatic division of existing naïve T cells.

Sustaining a highly diverse naïve T cell repertoire solely through replication of existing clones presents an interesting challenge. Naïve T cells compete for resources and space in tissues, which might lead to certain clones, particularly those made early in life, outcompeting unexpanded, newly produced clones later in life [33]. This amounts to a cellular equivalent of the ‘tragedy of the commons’, which posits that individuals acting in their own self-interest will deplete a shared resource, bringing about the eventual collapse of the system [54]. This is particularly true for access to IL-7 which is necessary for naïve T cell survival and proliferation, and is expressed at constant, low levels by stromal cells in the secondary lymphoid tissues [55]. The impact of IL-7 levels on naïve T cell turnover is confirmed by the observation that therapeutic administration of high levels of IL-7 promotes substantial increases in Ki67+ naïve T cells after 1 week [56].

This prompts the question: How do naïve T cells undergo peripheral expansion without leading to unequal clonal growth and collapse of diversity? The slow, constant turnover of the population, with estimated individual naïve T cell division rates on the order of once every 3-8 years, provides a general mechanism to support population diversity over 30-40 years after thymic involution. Additionally, we hypothesized that mechanisms are likely to exist to support equality in clonal growth across the diverse repertoire of naïve T cell clones and ensure that expanded clones do not consume space and resources at the expense of newly generated or unexpanded clones.

We provide evidence of such a mechanism, by monitoring cell age changes in the CD4^+^ naïve T cell population relative to subpopulations with different replicative histories. We observed that expanded populations of CD4^+^CD31^−^ naïve T cells are less likely to undergo homeostatic proliferation in the periphery as compared to less expanded CD4^+^CD31^+^ naïve T cells. Mechanistically, we show that this is partly regulated by changes in basal NF-κB activity that alter downstream effects of IL-7 stimulation, leading to restrictions on proliferation in response to the prototypical homeostatic growth factor important for naïve T cell survival and expansion. Such a mechanism could potentially favor proliferation of clones in the less expanded CD4^+^CD31^+^ compartment, facilitating even expansion of all naïve T cell clones in the periphery.

Interestingly, CD8^+^ naïve T cells appear to exhibit reduced homeostatic proliferation in vivo relative to CD4^+^ naïve T cells. However, *in vitro* we observed that CD8^+^ naïve T cells responded equivalently, if not more than, CD4^+^CD31^+^naïve T cells to IL-7, suggest that additional factors contribute to the regulated, slow rate of homeostatic division of CD8^+^ naïve T cells *in vivo.* A simple explanation would again be that IL-7 itself is highly limiting *in vivo* as exogenous IL-7 therapy led to a massive increase in Ki67^+^CD8^+^ naïve T cells (>50% of the population) within a week after administration[56].

An interesting implication of our findings is how inflammation could influence diversity within the naïve T cell pool. We demonstrate that addition of TNF, a factor commonly increased in inflammatory settings, allows for extensive proliferation of all naïve T cell subsets to IL-7, including expanded CD4^+^CD31^−^ naïve T cells which typically show minimal proliferation to IL-7 *in vitro.* Chronic inflammatory conditions such as rheumatoid arthritis are known to significantly alter the naïve T cell repertoire in humans [57, 58]. Moreover, the well described collapse in diversity seen in old age may be a direct consequence of increased chronic inflammation, termed ‘inflammaging,’ which is increasingly recognized as a major contributor to age-related immune dysfunction among other diseases [59, 60]. Given that there are approved therapies for TNF-inhibitors in the context of inflammatory disorders, it could be interesting to explore how these may influence naïve T cell diversity in elderly individuals [61]. Moreover, defining the factors that influence changes in naïve T cell diversity and the consequences of these changes in the elderly will be of increasing importance as human lifespans continue to increase in the coming decades.

## MATERIALS AND METHODS

### Isolation of Cells

Buffy coats were obtained from anonymous healthy blood donors at Blodcentralen, Karolinska University hospital with the approval of the ethical review board in Stockholm, Sweden (2006/229-31/3). Red blood cell lysis was performed by combining the total buffy coat with a (1:3) solution of RBC lysis buffer (155mM NH_4_CL, 12mM NaHCO_3_, 0.1mM EDTA) and incubated for 10 minutes at room temperature.

After the 10-minute incubation the samples were centrifuged in 50mL conical tubes and resuspended in 5ml PBS (containing 2mM EDTA) to which 45ml of RBC lysis buffer was again added followed immediately by a second centrifugation step. The remaining cells were washed two times in MACS buffer (PBS, 0.5% bovine serum albumin, 2mM EDTA) to remove residual lysis buffer. Samples were filtered through 40μM mesh filters.

### Magnetic Bead Purification

Total white blood cells were obtained by RBC lysis. An antibody cocktail designed for negative selection of purified naïve T cells (Naïve Pan T cell purification kit, Miltenyi Biotec) was used together with extra αCD15/CD14 beads. This allowed for a single selection step over D columns that gave highly purified CD4^+^ and CD8^+^ T naïve T cells (typically >90%) prior to fluorescence activated cell sorting (FACS).

### Flow Cytometry and Cell Sorting

Enriched populations of cells obtained after magnetic selection protocols were incubated with a cocktail of antibodies and sorted by FACS: CD3ε-FITC (clone UCHT1, BD Biosciences), CD4-APC Cy7 (clone RPA-T4, BD Biosciences), CD8α-BV768 (clone RPA-T8, BD Biosciences), CD45RA-PE-CF594 (clone HI100, BD Biosciences), CCR7-BV450 (clone 150503, BD Biosciences), and CD31-APC (clone AC128, Miltenyi Biotec). Cells were labeled in MACS buffer containing antibodies at 4°C for 30 minutes and washed extensively and filtered to remove dead cells and debris prior to sorting. FACS was performed using a BD Influx cell sorter equipped with blue (480nm), red (640nm), and violet (405nm) lasers. Sort purities were determined after sorting for all populations.

### Flow Cytometry Analysis of Ki67 Expression and Tscm Frequency

For analysis of Ki67 expression in T cell subsets, we thawed cryopreserved PBMC, and stained them with CD3ε-Alexa700 (clone UCHT1), CD4-APC-H7 (clone SK3), CCR7-BV421 (clone 150503), CD27-BV786 (clone L128), CD49F-BV650 (clone GoH3), (CD57-FITC (clone NK-1), CD14-V500 (clone MΦP9), CD19-V500 (clone HIB19) (all from BD Biosciences), CD127-PE-Cy7 (clone R34.34, Beckman Coulter), CD8a-BV570 (clone RPA-T8, BioLegend), CD95-PE (clone DX2, BioLegend), CD45RA-PE-Texas Red (clone MEM-56, ThermoFisher), CD31-APC (clone AC128, Miltenyi Biotec) and Live/Dead fixable Aqua dead cell stain (ThermoFisher) in PBS containing 2% fetal bovine serum and 2mM EDTA. After washing, the PBMCs were fixed with fixation/permeabilization buffer (Transcription Factor Staining Buffer, Ebioscience) for 30min at 4°C, washed twice with permeabilization buffer (Transcription Factor Staining Buffer, Ebioscience), and stained for 30min with anti-Ki67-FITC (clone 35/Ki-67, BD Biosciences) diluted in permeabilization buffer. Data was acquired on a BD Biosciences LSR Fortessa instrument equipped with 405nm, 488nm, 561nm, and 647nm lasers, and analyzed using FlowJo software (v9.7.8). Naïve CD4^+^ and CD8^+^ T cells were defined as CCR7^+^CD45RA^+^ CD95^−^ cells and stem cell memory T cells (T_scm_) were defined as CCR7^+^CD45RA^+^CD95^+^ cells, among CD3^+^CD4^+^ or CD3^+^CD8^+^ live, CD14^−^CD19^−^ lymphocytes.

### Analysis of phospho-NF-κB by flow cytometry

Peripheral blood was collected in heparin tubes and PBMCs were isolated using density centrifugation. The freshly isolated PBMC were stained with antibodies against CD31 Alexa 647(clone xx, company), CD14 V500 (clone MfP9, BD Biosciences), CD19 V500 (clone HIB19, BD Biosciences), CCR7 BV421 (clone 150503, BD Biosciences), CD8 BV570 (BioLegend), and LIVE/DEAD fixable dead cell stain near IR (Invitrogen) in RPMI1640 with 10% FCS (R10 medium) for 30 min at 4°C. After washing twice, 50ml (10^6^ PBMC) were added to 150ml prewarmed (37°C) media with or without 15ng/ml TNF, and incubated at 37°C for 3 min, after which 200 ml prewarmed 4% paraformaldehyde was added. The cells were fixed for 20 min at 37°C and for an additional 30 min at 4°C, washed twice in PBS containing 2mM EDTA and 1% FCS (FACS wash), and resuspended in 5ml FACS wash. Subsequently, 200ml methanol (−20°C) was added and the cells were incubated at −20°C for 3h, washed twice in FACS wash and incubated with the following antibodies for 1h at room temperature: CD4 FITC (clone SK3, BD Biosciences), CD3 Alexa 700 (clone UCHT1, BD Biosciences), CD45RA BV786 (clone HI100, BD Biosciences), and pNF-κB S529 PE-CF594 (clone K10-895.12.50, BD Biosciences). The cells were washed twice and immediately acquired on an LSR Fortessa (BD Biosciences), and analyzed using FlowJo v9.8 (Treestar Inc). Naïve CD4^+^ T cells were defined as CD45RA^+^CCR7^+^ among single live, CD14^−^CD19^−^CD3^+^CD4^+^ lymphocytes.

### DNA Isolation

The DNA extractions were done in an ISO8 cleanroom to reduce carbon contamination. Beforehand, the glass utensils were baked at 450°C for 4 hours. To each sample 1ml of lysis buffer (100 mM Tris [pH 8.0], 200 mM NaCl, 1% SDS, and 5 mM EDTA) and 12 μl of Proteinase K (40 mg/ml) were added. The sample were then incubated overnight at 65°C. 6μl of RNase cocktail (Ambion) was added to each sample and the samples were then incubated at 65°C for 1 hr. Each sample received half a volume of NaCl (5 M). After a 30 s vortex, the tubes were centrifuged at 13,000 rpm for 6 min. The pellet was discarded and the supernatant transferred to a glass tube with 3 volumes of absolute ethanol. The tubes were gently agitated for 30 seconds. The DNA pellet formed in the previous step was washed three times in buffer (70% Ethanol [v/v] and 0.5 M NaCl) and later dried at 65°C overnight to evaporate all the ethanol. 500 ml of DNase/RNAase free water (GIBCO/Invitrogen) was added to each vial and left overnight at 65°C to resuspended the DNA. The concentration and purity were assessed by UV spectroscopy (NanoDrop).

### Accelerator Mass Spectrometry

A special sample preparation method has been developed for the μg-sized DNA samples *(1)*. The purified DNA samples were received suspended in water. The samples were subsequently lyophilized to dryness under vacuum and centrifugation. Excess CuO was added to the dried samples in quartz tubes which were then evacuated and sealed with a high temperature torch. The quartz tubes were placed in a furnace set at 900°C for 3.5 hours to combust all carbon to CO_2_. The evolved gas was cryogenically purified and trapped. The CO_2_ gas was converted to graphite in individual sub-mL reactors at 550°C for 6 hours in the presence of zinc powder as reducing agent and iron powder as catalyst. The graphite targets were measured at the Department of Physics and Astronomy, Ion Physics, Uppsala University, using a 5 MV Pelletron tandem accelerator *(2)*. Stringent and thorough laboratory practice is necessary to minimize the introduction of stray carbon into the samples, including preheating of all glassware and chemicals prior to samples preparation. Large CO_2_ samples (>100 μg) were split and δ^13^C was measured by stable isotope ratio mass spectrometry, which established the δ^13^C correction to −24.1 ± 1 ‰ (2 SD) for leucocyte samples. Corrections and reduction of background contamination introduced during sample preparation were made as described by Hua et al *(3)* and Santos et al (4). The measurement error was determined for each sample and ranged between ±8 ‰ and 24 ‰ (2 SD) Δ^14^C for the large sample and small samples (10μgC) respectively. All ^14^C data are reported as decay-corrected Δ^14^C or Fraction Modern. All AMS analyses were performed blind to age and origin of the sample.

### Mass Cytometry and T cell activation assays

Total peripheral blood mononuclear cells (PBMC) were purified from buffy coats by density centrifugation (FH). PBMC were washed and subsequently re-suspended in cold cell culture medium (RPMI 10% FBS, L-Glu). The PBMCs were divided into separate tubes and incubated for 30 minutes on ice with biotinylated anti-CD3 (1μg/ml, OKT3; Miltenyi), anti-CD3/CD28 (CD28-10μg/ml, 15E8; Miltenyi)/CD2 (10μg/ml, LT2; Miltenyi), or nothing (non-stimulated control). All groups were incubated with anti-CD31 coupled to 148Nd prior to stimulation, for staining (not crosslinking) and initial experiments performed without pre-incubation of anti-CD31 gave comparable results for all measures, albeit with reduced resolution of CD31 expression (data not shown). The cells were washed one time with an excess of cell culture medium (100x), re-suspended in pre-warmed cell culture medium and transferred to 96-well U-bottom culture plates containing pre-diluted cytokines (IL-2 or IL-7 both at 10ng/ml final) and streptavidin (50μg /ml) to crosslink simulation antibodies. Individual reactions were terminated by the addition of a formaldehyde-based fixation buffer (Cytodelics, Stockholm Sweden) at different times after addition of cells and stored at 4°C prior to staining and mass cytometry analysis. Cells were washed to remove fixation buffer and re-suspended in staining buffer with antibodies detailed in Table S2. Cells were washed twice with staining buffer, filtered through a 35μm nylon mesh filter followed by dilution to 5×10^5^ cells/ml. Cells were analyzed on a CyTOF2 mass cytometer (Fluidigm, CA, USA) with software version 6.0.626 using a noise reduction with a lower convolution threshold of 200, a sigma value of 3 and an event length limitation of 10-150. FCS 3.0 files were exported without randomization and then normalized using internal bead-standards (5). Cells were gated on as having an event length between 20-65 and being positive for Iridium DNA-intercalator. Naïve T cell subsets were defined as lineage negative (CD11c, CD19, CD235a/b) and CD3^+^CD45RA^+^CCR7^+^CD27^+^ with CD4^+^ or CD8^+^ cells gated separately. Events were exported for down-stream ACCENSE and DREVI analyses. The single-cell data was visualized by ACCENSE with CD4^+^CD31^+^ and CD31^−^ naïve as well as memory/effector CD4^+^ T cells combined for dimensionality reduction. DREVI was performed on manually gated cell populations with naïve defined as above, and memory/effector cells defined as all non-naïve T cells within the CD4^+^ T cell fraction.

### *In vitro* T cell proliferation assays

Naïve T cells were purified as described for ^14^C dating protocols. The cells were subsequently washed in PBS and labeled with 5μM Carboxyfluorescein succinimidyl ester ((CFSE) CellTrace, Invitrogen) in PBS for 10 minutes at 37°C and washed extensively in cell culture medium. Labeled cells were added to 96-well U-bottom tissue culture plates. Specified wells were pre-coated with αCD3 (HIT3A, BD Biosciences) by incubation with PBS containing anti-CD3 (0.1μg/ml) at 37°C for 4 hours and extensive washing to remove unbound antibody. Unstimulated cells were added to wells that had not been pre-coated with anti-CD3. Homeostatic proliferation assays were performed by culturing cells with recombinant human IL-7 (RND systems) at 10ng/ml added on Days 0, 2, 4, and 6 in the presence or absence of anti-CD3. Activation assays were performed by culturing cells with recombinant human IL-2 (10ng/ml, RND Systems) in the presence of coated anti-CD3 and anti-CD28 (2mg/ml, BD Biosciences) in the culture medium. For NF-κB inhibition assays, the inhibitor 6-amino-4–4-phenoxyphenylethylamino-quinazoline (Merck Millipore, USA) was reconstituted in DMSO at a stock concentration of 1mM and administered at a final concentration of 10nM on Days 0 and 4. This concentration was chosen based on published ranges of toxicity for human T cells [49]. Control wells not receiving inhibitor, received an equal volume of DMSO (1:100,000 dilution in culture medium) at the time inhibitor was added. Proliferation was measured on day 8 by labeling cells with standard memory/naïve phenotyping antibodies used for naïve purification (excluding CD3-FITC due to CFSE labeling) and performing flow cytometry analysis to measure CFSE dilution within naïve (or memory) T cell subsets (analysis done using FlowJo software version 9.2).

### Mathematical Modeling

Statistical and mathematical analyses of data relating to ^14^C measurements are included as an attached SSM.

## CONTRIBUTIONS

Contributed to study design: JF, PR, JEM, JM, SB. Performed isolation of cells and gDNA for ^14^C measurements: PR, JEM, AK. Designed mass cytometry studies and performed analysis of mass cytometry data: PR, JEM, AO, PB. Performed multi-parameter flow cytometry for phenotyping and Ki67/NF-κB studies by FACS: JEM, JM. Performed all mathematical modeling and computer simulations: SB. Performed ^14^C measurements on DNA and interpreted AMS results: MS, GP. Provided conceptual advice and assisted with manuscript preparation: SR and AY. Prepared figures and wrote manuscript: PR, JEM, SB, AO, PB, JM and JF.

## ACKNOWLEDGMENTS

We are grateful to Marcelo Toro and Sarantis Giatrellis for help with flow cytometry and Karl Håkansson and Peter Senneryd for AMS sample preparation. This study was supported by grants from the Swedish Research Council, the Swedish Cancer Society, the Karolinska Institute, the Strategic Research Programme in Stem Cells and Regenerative Medicine at Karolinska Institutet (StratRegen), Knut och Alice Wallenbergs Stiftelse and Torsten Söderbergs Stiftelse. Pedro Réu was supported by the Portuguese Foundation for Science and Technology (SFRH/BD/33465/2008). Jeff E. Mold was supported by a Human Frontiers Science Program Long-Term Fellowship (LT-000231/2011-L). Petter Brodin was supported by the Swedish Society for Medical Research.

## SUPPLEMENTARY INFORMATION

### Supplemental Figures

**S Fig 1:**
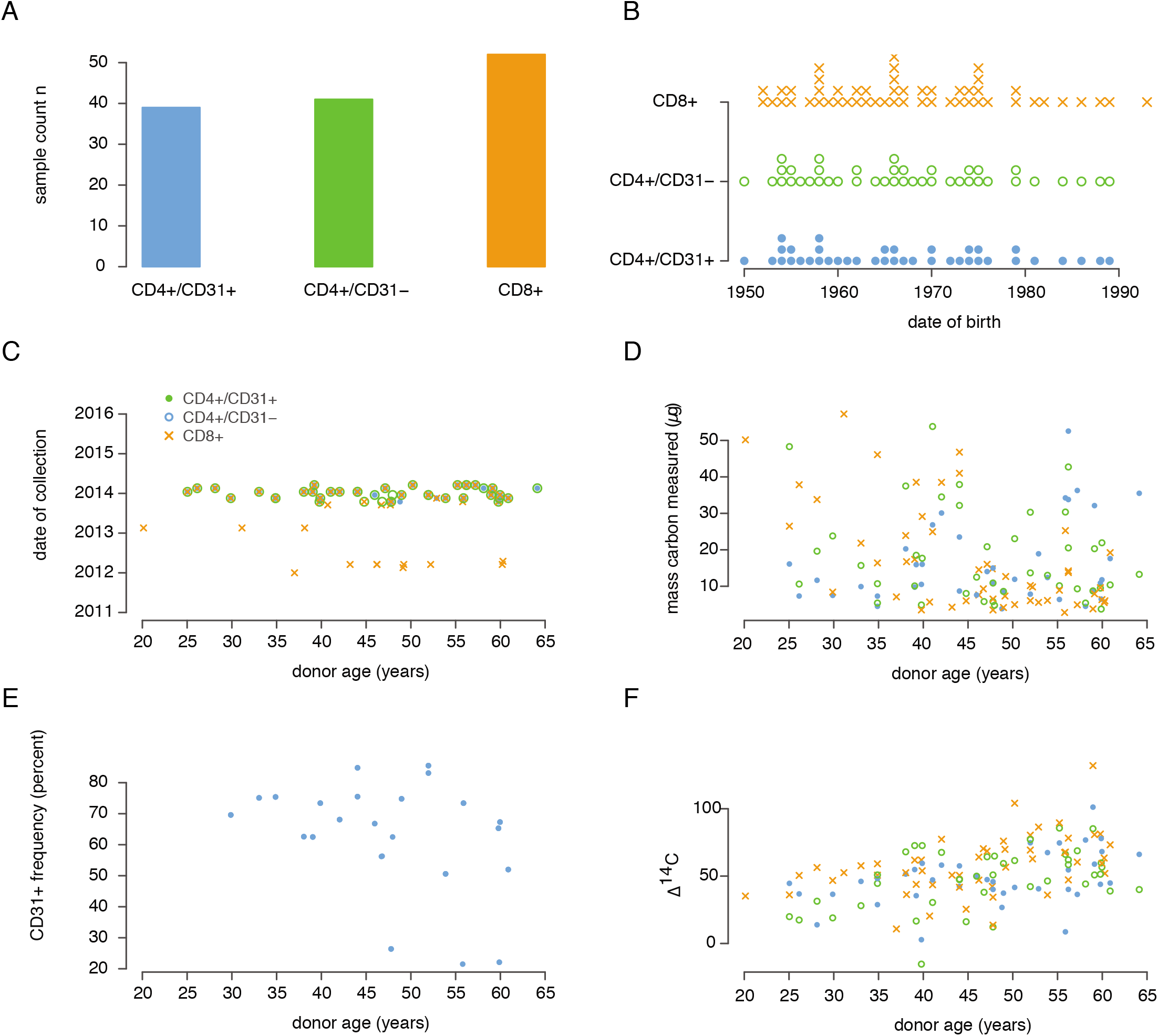
Summary of Donor Characteristics and Samples Involved in ^14^C Study. A complete overview is also provided in Supplemental Table 1. (A) Total number of samples included in ^14^C cell age determination for Figures 1–4. (B) Distribution of samples relative to donor age (date of birth) for each sample set. (C) Collection date for each sample relative to donor age. Most samples were collected in 2014. (D) Carbon mass after isolation from purified DNA taken from each sample relative to donor age. (E) Frequency of CD31^+^ cells within the CD4^+^ naïve T cell fraction for all donors. (F) Summary of ^14^C measurements for all donors and cell types in study.

**S Fig 2:**
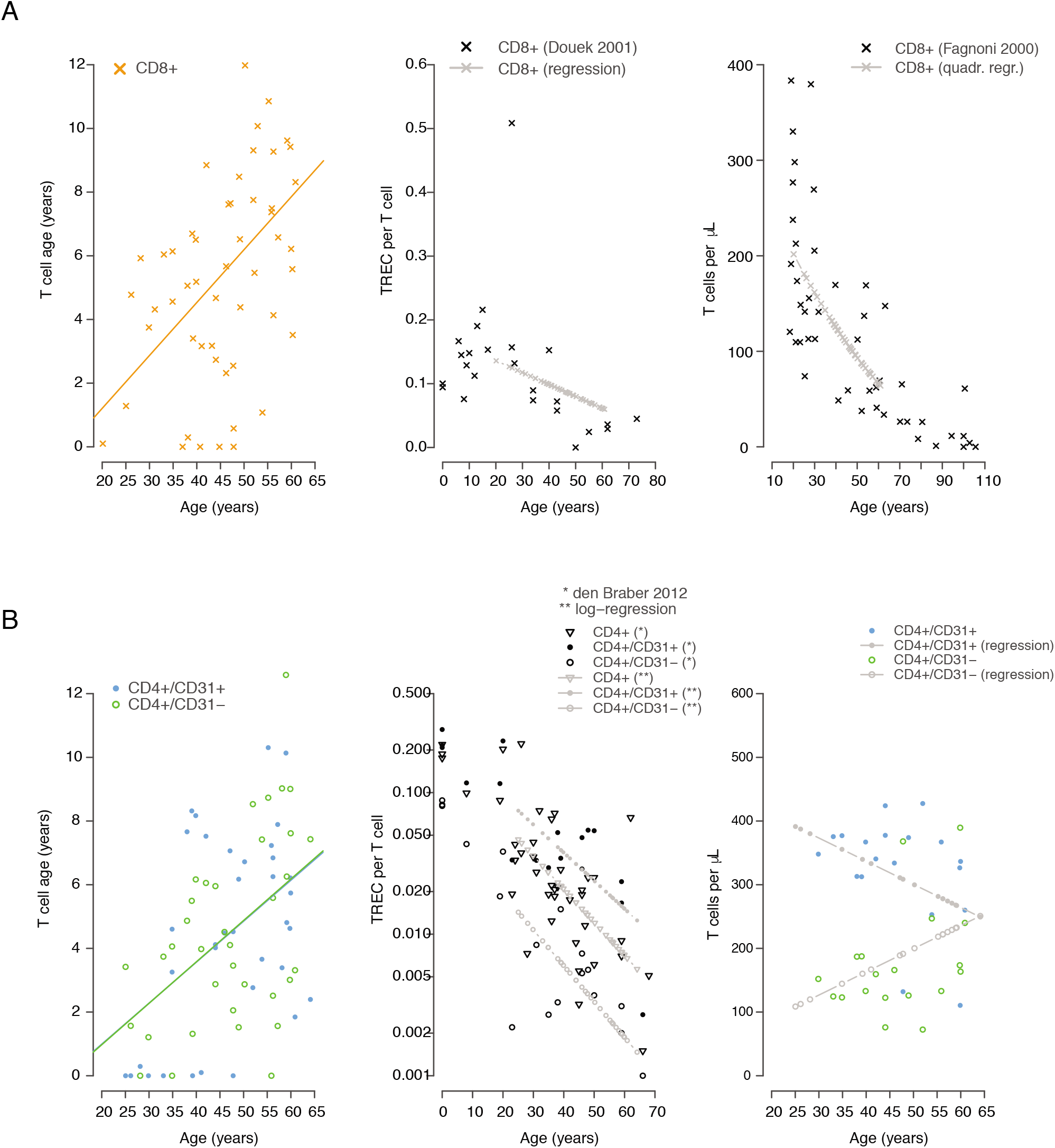
Regressed data used for linear models presented used in Figures 3–4 to define dynamics of naïve T cell homeostasis. (A) CD8^+^ naïve T cell measurements for cell age (based on this study), TREC content (taken from ref: [23]), and cell numbers (taken from ref: [13]). (B) CD4^+^ naïve T cell measurements for cell age (based on this study), TREC content (taken from ref: [8]), and cell numbers. Cell numbers were set to 500 cells/μl and defined in terms of CD31^+^ and CD31-fractions based on regressed data from S Fig 1E.

**S Fig 3:**
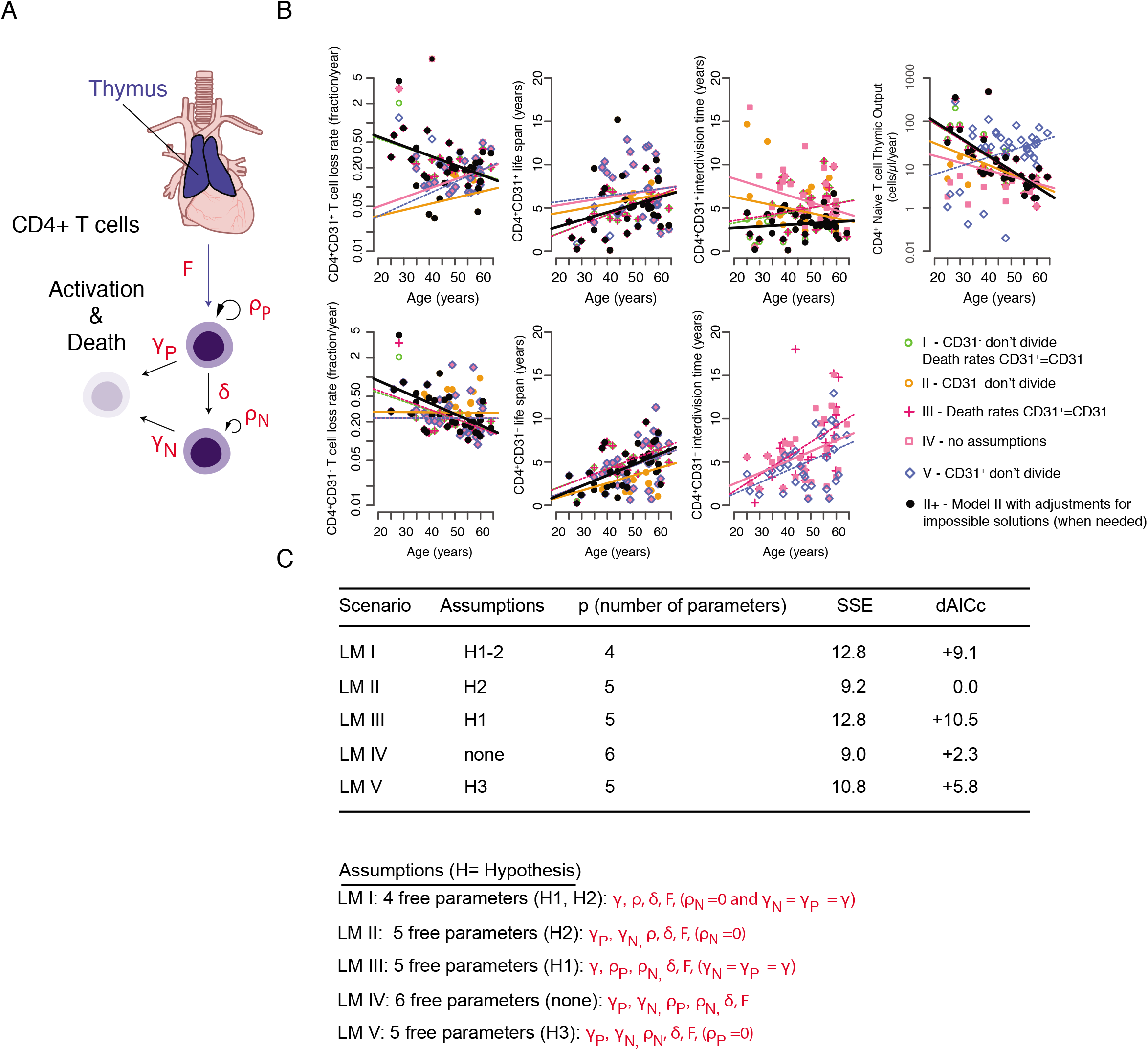
Hypothesis testing for CD4^+^ naïve T cell dynamics using linear models relevant to Figure 3. (A) Schematic of CD4^+^ naïve T cell production, proliferation, differentiation and activation/death. (B) Representation of calculated dynamic values for each scenario tested. (C) Table indicating different features of each scenario (Linear Models I-V) and sum of squared errors (SSE) and ΔAICc (dAICc) for each scenario. Hypotheses tested for each scenario are listed below.

**S Fig 4:**
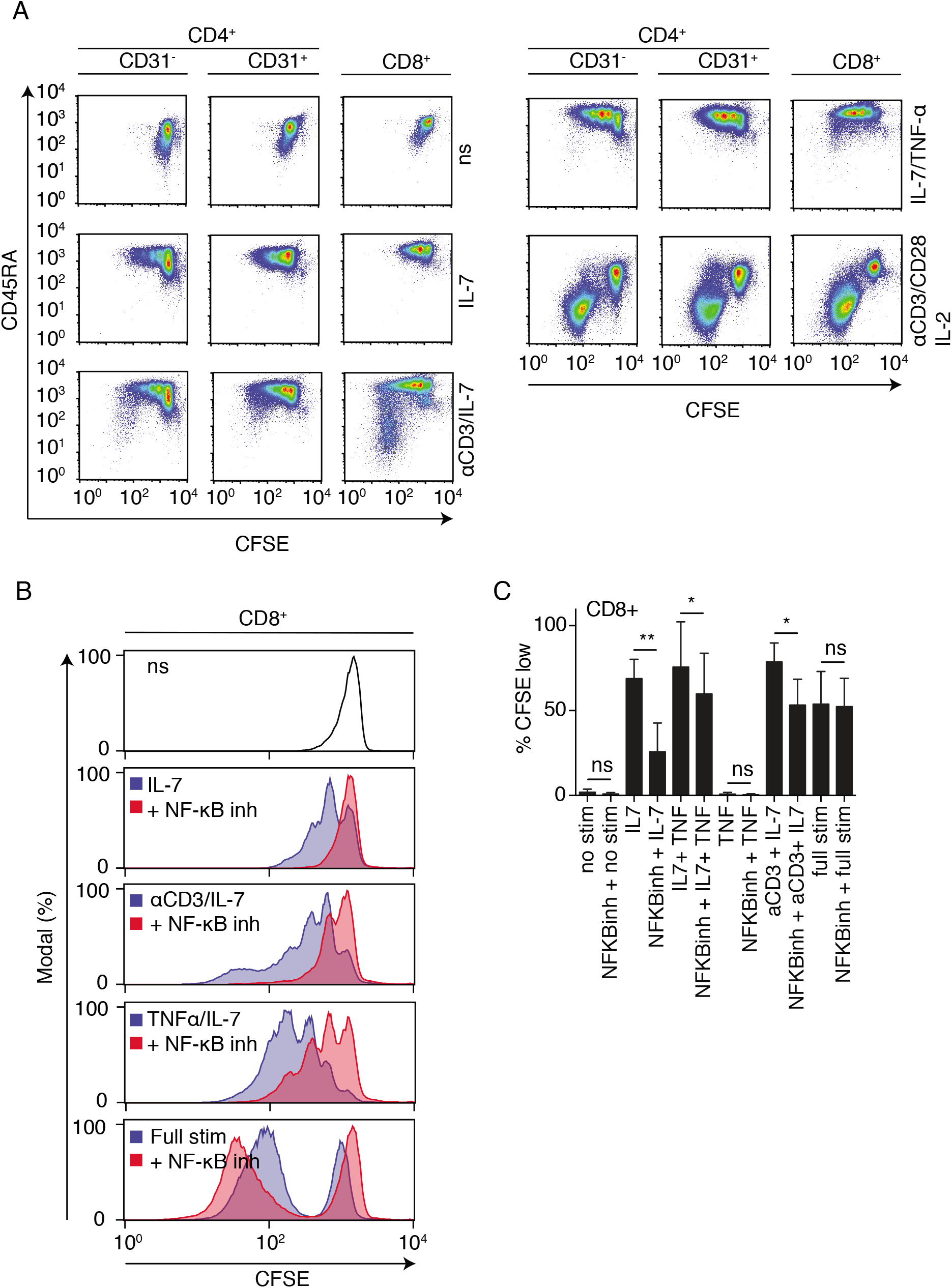
Monitoring activation status of in vitro stimulated CD4^+^ and CD8^+^ naïve T cells in CFSE assays from Figure 6. (A) CFSE dilution versus CD45RA expression for each stimulation condition. Minimal CD45RA downregulation is observed with the exception aCD3 ^+^ IL-7 stimulation. Full stimulation (αCD3/αCD28 ^+^ IL-2) results in full activation of naïve T cells. (B) CFSE dilution with different conditions for CD8^+^ naïve T cells. Vehicle (DMSO – blue histograms) and treatment with NF-κB inhibitor (blue histogram) is shown. (C) Summary of four independent donors for CD8^+^ naïve T cells.

**S Fig 5:**
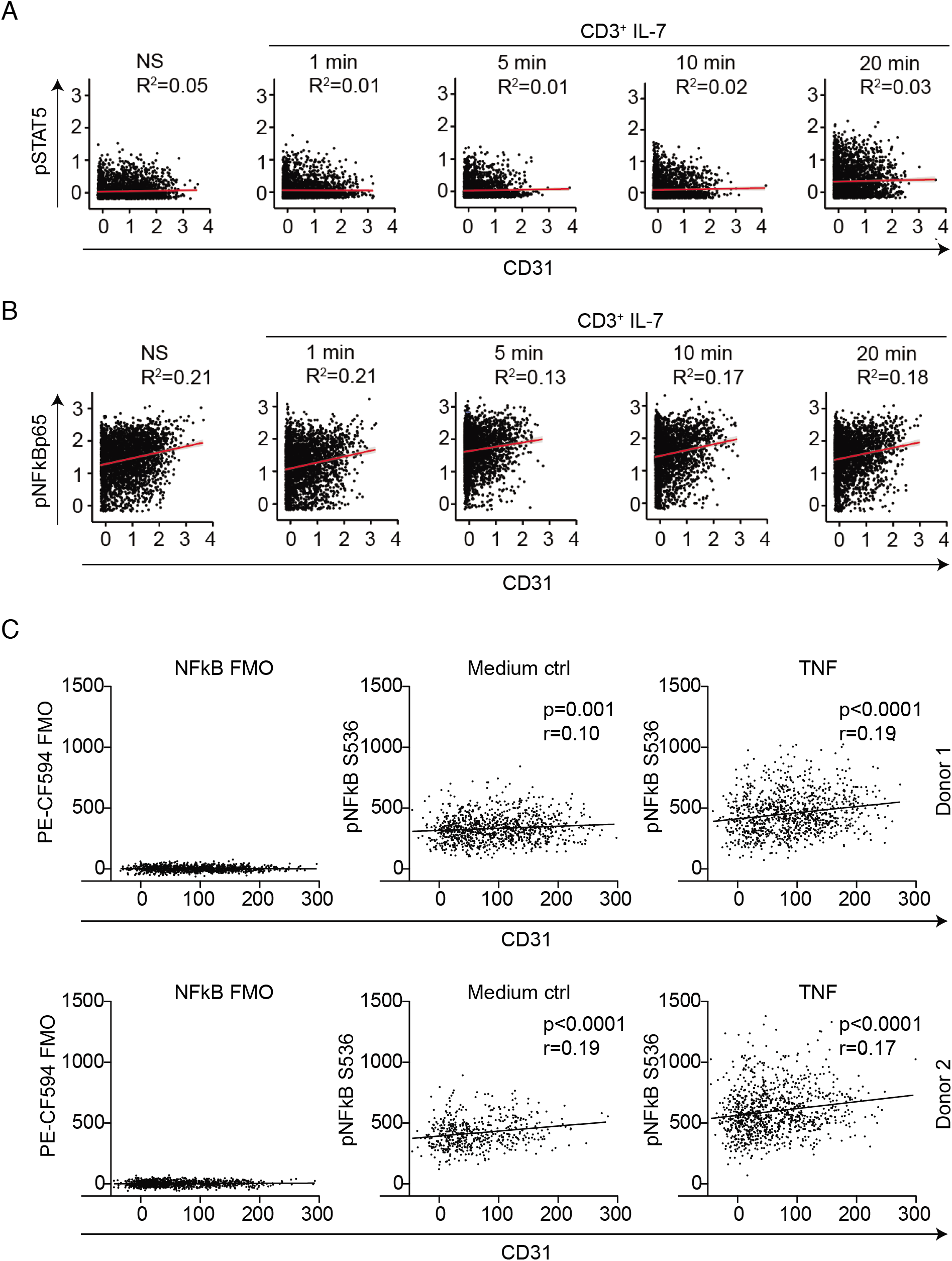
Extended data for Figure 6 showing raw correlations between CD31 and phosphor-proteins from CyTOF data and confirmation using Flow Cytometry. (A) Phosphorylation of STAT5 in unstimulated (NS) and in aCD3 ^+^ IL-7 (10ng/mL) stimulated peripheral blood mononuclear cells. CD4^+^ Naïve T cells are identified by gating on lineage negative, CD3^+^CD4^+^CD45RA^+^CD27^+^CCR7^+^ and Pearson’s correlations are depicted for phosphor-STAT5 (y-axis) versus CD31 expression (x-axis). Values for different timepoints post-stimulation are shown (B) Phosphor-NF-κB (RelA/p65) versus CD31 expression on the same populations as in (A). (C) Confirmation of CyTOF results using flow cytometry to identify naïve T cells (gate: Live, Lineage Negative, CD3^+^CD4^+^CD45RA^+^CCR7^+^) and monitoring phosphor-NF-κB (RelA/p65) (y-axis) versus CD31 expression (x-axis) in unstimulated (middle panels) and TNF stimulated (right panels) PBMCs. Top and bottom panels represent two different healthy adult donors. Background fluorescence for phosphor-NF-κB is shown in the left panels (Fluorescence Minus One (FMO)).

**S Table 1**: Summary of all donors and measurements for ^14^C testing.

**S Table 2**: Scenario testing for Figure 2 to define best fit scenarios for proportion of naïve T cells that undergo turnover in the periphery.

**S Table 3**: Robustness tests for all modeling results for Figures 1–4.

**S Table 4**: List of antibodies used for mass cytometry studies in Figure 6.

## REFERENCES

1. van den Broek T, Borghans JAM, van Wijk F. The full spectrum of human naive T cells. Nat Rev Immunol. 2018;18(6):363–73. Epub 2018/03/10. doi: 10.1038/s41577-018-0001-y. PubMed PMID: 29520044.

2. Nikolich-Zugich J, Slifka MK, Messaoudi I. The many important facets of T-cell repertoire diversity. Nat Rev Immunol. 2004;4(2):123–32. doi: 10.1038/nri1292. PubMed PMID: 15040585.

3. von Gaudecker B. Ultrastructure of the age-involuted adult human thymus. Cell Tissue Res. 1978;186(3):507–25. PubMed PMID: 304768.

4. Steinmann GG, Klaus B, Muller-Hermelink HK. The involution of the ageing human thymic epithelium is independent of puberty. A morphometric study. Scand J Immunol. 1985;22(5):563–75. PubMed PMID: 4081647.

5. Bains I, Antia R, Callard R, Yates AJ. Quantifying the development of the peripheral naive CD4+ T-cell pool in humans. Blood. 2009;113(22):5480–7. doi: 10.1182/blood-2008-10-184184. PubMed PMID: 19179300; PubMed Central PMCID: PMC2689049.

6. McDade TW, Worthman CM. Evolutionary process and the ecology of human immune function. Am J Hum Biol. 1999;11(6):705–17. doi: 10.1002/(SICI)1520-6300(199911/12)11:6<705::AID-AJHB1>3.0.CO;2-G. PubMed PMID: 11533987.

7. Flores KG, Li J, Sempowski GD, Haynes BF, Hale LP. Analysis of the human thymic perivascular space during aging. J Clin Invest. 1999;104(8):1031–9. doi: 10.1172/JCI7558. PubMed PMID: 10525041; PubMed Central PMCID: PMC408578.

8. den Braber I, Mugwagwa T, Vrisekoop N, Westera L, Mogling R, de Boer AB, et al. Maintenance of peripheral naive T cells is sustained by thymus output in mice but not humans. Immunity. 2012;36(2):288–97. doi: 10.1016/j.immuni.2012.02.006. PubMed PMID: 22365666.

9. Surh CD, Sprent J. Homeostasis of naive and memory T cells. Immunity. 2008;29(6):848–62. doi: 10.1016/j.immuni.2008.11.002. PubMed PMID: 19100699.

10. Johnson PL, Yates AJ, Goronzy JJ, Antia R. Peripheral selection rather than thymic involution explains sudden contraction in naive CD4 T-cell diversity with age. Proc Natl Acad Sci U S A. 2012;109(52):21432–7. Epub 2012/12/14. doi: 10.1073/pnas.1209283110. PubMed PMID: 23236163; PubMed Central PMCID: PMCPMC3535632.

11. Qi Q, Liu Y, Cheng Y, Glanville J, Zhang D, Lee JY, et al. Diversity and clonal selection in the human T-cell repertoire. Proc Natl Acad Sci U S A. 2014;111(36):13139–44. doi: 10.1073/pnas.1409155111. PubMed PMID: 25157137; PubMed Central PMCID: PMC4246948.

12. Goronzy JJ, Fang F, Cavanagh MM, Qi Q, Weyand CM. Naive T cell maintenance and function in human aging. J Immunol. 2015;194(9):4073–80. doi: 10.4049/jimmunol.1500046. PubMed PMID: 25888703; PubMed Central PMCID: PMC4452284.

13. Fagnoni FF, Vescovini R, Passeri G, Bologna G, Pedrazzoni M, Lavagetto G, et al. Shortage of circulating naive CD8(+) T cells provides new insights on immunodeficiency in aging. Blood. 2000;95(9):2860–8. PubMed PMID: 10779432.

14. Vrisekoop N, den Braber I, de Boer AB, Ruiter AF, Ackermans MT, van der Crabben SN, et al. Sparse production but preferential incorporation of recently produced naive T cells in the human peripheral pool. Proc Natl Acad Sci U S A. 2008;105(16):6115–20. doi: 10.1073/pnas.0709713105. PubMed PMID: 18420820; PubMed Central PMCID: PMC2329696.

15. De Boer RJ, Perelson AS. Quantifying T lymphocyte turnover. J Theor Biol. 2013;327:45–87. doi: 10.1016/j.jtbi.2012.12.025. PubMed PMID: WOS:000318258400005.

16. Westera L, van Hoeven V, Drylewicz J, Spierenburg G, van Velzen JF, de Boer RJ, et al. Lymphocyte maintenance during healthy aging requires no substantial alterations in cellular turnover. Aging Cell. 2015;14(2):219–27. Epub 2015/01/30. doi: 10.1111/acel.12311. PubMed PMID: 25627171; PubMed Central PMCID: PMCPMC4364834.

17. Gattinoni L, Lugli E, Ji Y, Pos Z, Paulos CM, Quigley MF, et al. A human memory T cell subset with stem cell-like properties. Nat Med. 2011;17(10):1290–7. doi: 10.1038/nm.2446. PubMed PMID: 21926977; PubMed Central PMCID: PMC3192229.

18. Kimmig S, Przybylski GK, Schmidt CA, Laurisch K, Mowes B, Radbruch A, et al. Two subsets of naive T helper cells with distinct T cell receptor excision circle content in human adult peripheral blood. J Exp Med. 2002;195(6):789–94. PubMed PMID: 11901204; PubMed Central PMCID: PMC2193736.

19. McFarland RD, Douek DC, Koup RA, Picker LJ. Identification of a human recent thymic emigrant phenotype. Proc Natl Acad Sci U S A. 2000;97(8):4215–20. Epub 2000/03/29. doi: 10.1073/pnas.070061597. PubMed PMID: 10737767; PubMed Central PMCID: PMCPMC18202.

20. Hazenberg MD, Borghans JA, de Boer RJ, Miedema F. Thymic output: a bad TREC record. Nat Immunol. 2003;4(2):97–9. doi: 10.1038/ni0203-97. PubMed PMID: 12555089.

21. Hazenberg MD, Otto SA, Cohen Stuart JW, Verschuren MC, Borleffs JC, Boucher CA, et al. Increased cell division but not thymic dysfunction rapidly affects the T-cell receptor excision circle content of the naive T cell population in HIV-1 infection. Nat Med. 2000;6(9):1036–42. doi: 10.1038/79549. PubMed PMID: 10973325.

22. Hazenberg MD, Verschuren MC, Hamann D, Miedema F, van Dongen JJ. T cell receptor excision circles as markers for recent thymic emigrants: basic aspects, technical approach, and guidelines for interpretation. J Mol Med (Berl). 2001;79(11):631–40. doi: 10.1007/s001090100271. PubMed PMID: 11715066.

23. Douek DC, Betts MR, Hill BJ, Little SJ, Lempicki R, Metcalf JA, et al. Evidence for increased T cell turnover and decreased thymic output in HIV infection. J Immunol. 2001;167(11):6663–8. PubMed PMID: 11714838.

24. Douek DC, McFarland RD, Keiser PH, Gage EA, Massey JM, Haynes BF, et al. Changes in thymic function with age and during the treatment of HIV infection. Nature. 1998;396(6712):690–5. doi: 10.1038/25374. PubMed PMID: 9872319.

25. Poulin JF, Viswanathan MN, Harris JM, Komanduri KV, Wieder E, Ringuette N, et al. Direct evidence for thymic function in adult humans. J Exp Med. 1999;190(4):479–86. PubMed PMID: 10449519; PubMed Central PMCID: PMC2195604.

26. Spalding KL, Bhardwaj RD, Buchholz BA, Druid H, Frisen J. Retrospective birth dating of cells in humans. Cell. 2005;122(1):133–43. doi: 10.1016/j.cell.2005.04.028. PubMed PMID: 16009139.

27. Levin I, Kromer B. The tropospheric (CO2)-C-14 level in mid-latitudes of the Northern Hemisphere (1959-2003). Radiocarbon. 2004;46(3):1261–72. PubMed PMID: WOS:000231338100017.

28. Spalding KL, Arner E, Westermark PO, Bernard S, Buchholz BA, Bergmann O, et al. Dynamics of fat cell turnover in humans. Nature. 2008;453(7196):783–7. doi: 10.1038/nature06902. PubMed PMID: 18454136.

29. Yeung MS, Zdunek S, Bergmann O, Bernard S, Salehpour M, Alkass K, et al. Dynamics of oligodendrocyte generation and myelination in the human brain. Cell. 2014;159(4):766–74. doi: 10.1016/j.cell.2014.10.011. PubMed PMID: 25417154.

30. Bergmann O, Bhardwaj RD, Bernard S, Zdunek S, Barnabe-Heider F, Walsh S, et al. Evidence for cardiomyocyte renewal in humans. Science. 2009;324(5923):98–102. doi: 10.1126/science.1164680. PubMed PMID: 19342590; PubMed Central PMCID: PMCPMC2991140.

31. Landsverk OJ, Snir O, Casado RB, Richter L, Mold JE, Reu P, et al. Antibody-secreting plasma cells persist for decades in human intestine. J Exp Med. 2017. doi: 10.1084/jem.20161590. PubMed PMID: 28104812.

32. Rane S, Hogan T, Seddon B, Yates AJ. Age is not just a number: Naive T cells increase their ability to persist in the circulation over time. PLoS Biol. 2018;16(4):e2003949. Epub 2018/04/12. doi: 10.1371/journal.pbio.2003949. PubMed PMID: 29641514; PubMed Central PMCID: PMCPMC5894957.

33. Takada K, Jameson SC. Naive T cell homeostasis: from awareness of space to a sense of place. Nat Rev Immunol. 2009;9(12):823–32. doi: 10.1038/nri2657. PubMed PMID: 19935802.

34. Hataye J, Moon JJ, Khoruts A, Reilly C, Jenkins MK. Naive and memory CD4+ T cell survival controlled by clonal abundance. Science. 2006;312(5770):114–6. doi: 10.1126/science.1124228. PubMed PMID: 16513943.

35. Bains I, Thiebaut R, Yates AJ, Callard R. Quantifying thymic export: combining models of naive T cell proliferation and TCR excision circle dynamics gives an explicit measure of thymic output. J Immunol. 2009;183(7):4329–36. doi: 10.4049/jimmunol.0900743. PubMed PMID: 19734223.

36. Kohler S, Thiel A. Life after the thymus: CD31+ and CD31-human naive CD4+ T-cell subsets. Blood. 2009;113(4):769–74. doi: 10.1182/blood-2008-02-139154. PubMed PMID: 18583570.

37. Marelli-Berg FM, Clement M, Mauro C, Caligiuri G. An immunologist’s guide to CD31 function in T-cells. J Cell Sci. 2013;126(Pt 11):2343–52. Epub 2013/06/14. doi: 10.1242/jcs.124099. PubMed PMID: 23761922.

38. Kilpatrick RD, Rickabaugh T, Hultin LE, Hultin P, Hausner MA, Detels R, et al. Homeostasis of the naive CD4+ T cell compartment during aging. J Immunol. 2008;180(3):1499–507. PubMed PMID: 18209045; PubMed Central PMCID: PMC2940825.

39. Kohler S, Wagner U, Pierer M, Kimmig S, Oppmann B, Mowes B, et al. Post-thymic in vivo proliferation of naive CD4+ T cells constrains the TCR repertoire in healthy human adults. Eur J Immunol. 2005;35(6):1987–94. Epub 2005/05/24. doi: 10.1002/eji.200526181. PubMed PMID: 15909312.

40. Akaike H. New Look at Statistical-Model Identification. Ieee T Automat Contr. 1974;Ac19(6):716–23. doi: Doi 10.1109/Tac.1974.1100705. PubMed PMID: WOS:A1974U921700011.

41. Azevedo RI, Soares MV, Barata JT, Tendeiro R, Serra-Caetano A, Victorino RM, et al. IL-7 sustains CD31 expression in human naive CD4+ T cells and preferentially expands the CD31+ subset in a PI3K-dependent manner. Blood. 2009;113(13):2999–3007. doi: 10.1182/blood-2008-07-166223. PubMed PMID: 19008454.

42. Silva SL, Albuquerque AS, Matoso P, Charmeteau-de-Muylder B, Cheynier R, Ligeiro D, et al. IL-7-Induced Proliferation of Human Naive CD4 T-Cells Relies on Continued Thymic Activity. Front Immunol. 2017;8:20. Epub 2017/02/06. doi: 10.3389/fimmu.2017.00020. PubMed PMID: 28154568; PubMed Central PMCID: PMCPMC5243809.

43. Scholzen T, Gerdes J. The Ki-67 protein: from the known and the unknown. J Cell Physiol. 2000;182(3):311–22. doi: 10.1002/(SICI)1097-4652(200003)182:3<311::AID-JCP1>3.0.CO;2-9. PubMed PMID: 10653597.

44. van der Maaten L, Hinton G. Visualizing Data using t-SNE. J Mach Learn Res. 2008;9:2579–605. PubMed PMID: WOS:000262637600007.

45. Shekhar K, Brodin P, Davis MM, Chakraborty AK. Automatic Classification of Cellular Expression by Nonlinear Stochastic Embedding (ACCENSE). Proc Natl Acad Sci U S A. 2014;111(1):202–7. doi: 10.1073/pnas.1321405111. PubMed PMID: 24344260; PubMed Central PMCID: PMCPMC3890841.

46. Krishnaswamy S, Spitzer MH, Mingueneau M, Bendall SC, Litvin O, Stone E, et al. Systems biology. Conditional density-based analysis of T cell signaling in single-cell data. Science. 2014;346(6213):1250689. doi: 10.1126/science.1250689. PubMed PMID: 25342659; PubMed Central PMCID: PMCPMC4334155.

47. Kieper WC, Burghardt JT, Surh CD. A role for TCR affinity in regulating naive T cell homeostasis. J Immunol. 2004;172(1):40–4. Epub 2003/12/23. PubMed PMID: 14688307.

48. Silva A, Cornish G, Ley SC, Seddon B. NF-kappaB signaling mediates homeostatic maturation of new T cells. Proc Natl Acad Sci U S A. 2014;111(9):E846–55. doi: 10.1073/pnas.1319397111. PubMed PMID: 24550492; PubMed Central PMCID: PMC3948292.

49. Miller ML, Mashayekhi M, Chen L, Zhou P, Liu X, Michelotti M, et al. Basal NF-kappaB controls IL-7 responsiveness of quiescent naive T cells. Proc Natl Acad Sci U S A. 2014;111(20):7397–402. doi: 10.1073/pnas.1315398111. PubMed PMID: 24799710; PubMed Central PMCID: PMC4034246.

50. Goronzy JJ, Weyand CM. Successful and Maladaptive T Cell Aging. Immunity. 2017;46(3):364–78. Epub 2017/03/23. doi: 10.1016/j.immuni.2017.03.010. PubMed PMID: 28329703; PubMed Central PMCID: PMCPMC5433436.

51. Pulko V, Davies JS, Martinez C, Lanteri MC, Busch MP, Diamond MS, et al. Human memory T cells with a naive phenotype accumulate with aging and respond to persistent viruses. Nat Immunol. 2016;17(8):966–75. Epub 2016/06/09. doi: 10.1038/ni.3483. PubMed PMID: 27270402; PubMed Central PMCID: PMCPMC4955715.

52. Hale JS, Boursalian TE, Turk GL, Fink PJ. Thymic output in aged mice. Proc Natl Acad Sci U S A. 2006;103(22):8447–52. Epub 2006/05/24. doi: 10.1073/pnas.0601040103. PubMed PMID: 16717190; PubMed Central PMCID: PMCPMC1482512.

53. George AJ, Ritter MA. Thymic involution with ageing: obsolescence or good housekeeping? Immunol Today. 1996;17(6):267–72. Epub 1996/06/01. PubMed PMID: 8962629.

54. Hardin G. The tragedy of the commons. The population problem has no technical solution; it requires a fundamental extension in morality. Science. 1968;162(3859):1243–8. PubMed PMID: 5699198.

55. Link A, Vogt TK, Favre S, Britschgi MR, Acha-Orbea H, Hinz B, et al. Fibroblastic reticular cells in lymph nodes regulate the homeostasis of naive T cells. Nat Immunol. 2007;8(11):1255–65. Epub 2007/09/26. doi: 10.1038/ni1513. PubMed PMID: 17893676.

56. Sportes C, Hakim FT, Memon SA, Zhang H, Chua KS, Brown MR, et al. Administration of rhIL-7 in humans increases in vivo TCR repertoire diversity by preferential expansion of naive T cell subsets. J Exp Med. 2008;205(7):1701–14. Epub 2008/06/25. doi: 10.1084/jem.20071681. PubMed PMID: 18573906; PubMed Central PMCID: PMCPMC2442646.

57. Koetz K, Bryl E, Spickschen K, O’Fallon WM, Goronzy JJ, Weyand CM. T cell homeostasis in patients with rheumatoid arthritis. Proc Natl Acad Sci U S A. 2000;97(16):9203–8. Epub 2000/08/02. PubMed PMID: 10922071; PubMed Central PMCID: PMCPMC16846.

58. Wagner UG, Koetz K, Weyand CM, Goronzy JJ. Perturbation of the T cell repertoire in rheumatoid arthritis. Proc Natl Acad Sci U S A. 1998;95(24):14447–52. Epub 1998/11/25. PubMed PMID: 9826720; PubMed Central PMCID: PMCPMC24393.

59. Fulop T, Larbi A, Dupuis G, Le Page A, Frost EH, Cohen AA, et al. Immunosenescence and Inflamm-Aging As Two Sides of the Same Coin: Friends or Foes? Front Immunol. 2017;8:1960. Epub 2018/01/30. doi: 10.3389/fimmu.2017.01960. PubMed PMID: 29375577; PubMed Central PMCID: PMCPMC5767595.

60. Franceschi C, Bonafe M, Valensin S, Olivieri F, De Luca M, Ottaviani E, et al. Inflamm-aging. An evolutionary perspective on immunosenescence. Ann N Y Acad Sci. 2000;908:244–54.Epub 2000/07/27. PubMed PMID: 10911963.

61. Monaco C, Nanchahal J, Taylor P, Feldmann M. Anti-TNF therapy: past, present and future. Int Immunol. 2015;27(1):55–62. Epub 2014/11/21. doi: 10.1093/intimm/dxu102. PubMed PMID: 25411043; PubMed Central PMCID: PMCPMC4279876.

